# Enhanced RNA-targeting CRISPR-Cas technology in zebrafish

**DOI:** 10.1101/2024.10.08.617220

**Authors:** Ismael Moreno-Sanchez, Luis Hernandez-Huertas, Daniel Nahon-Cano, Carlos Gomez-Marin, Pedro Manuel Martinez-García, Anthony J. Treichel, Laura Tomas-Gallardo, Gabriel da Silva Pescador, Gopal Kushawah, Alejandro Díaz-Moscoso, Alejandra Cano-Ruiz, John A. Walker, Manuel J. Muñoz, Kevin Holden, Joan Galcerán, María Ángela Nieto, Ariel Bazzini, Miguel A. Moreno-Mateos

## Abstract

CRISPR-Cas13 systems are widely used in basic and applied sciences. However, its application has recently generated controversy due to collateral activity in mammalian cells and mouse models. Moreover, its efficiency could be improved in vivo. Here, we optimized transient formulations as ribonucleoprotein complexes or mRNA-gRNA combinations to enhance the CRISPR-RfxCas13d system in zebrafish. We i) used chemically modified gRNAs to allow more penetrant loss-of-function phenotypes, ii) improved nuclear RNA-targeting, and iii) compared different computational models and determined the most accurate to predict gRNA activity in vivo. Furthermore, we demonstrated that transient CRISPR-RfxCas13d can effectively deplete endogenous mRNAs in zebrafish embryos without inducing collateral effects, except when targeting extremely abundant and ectopic RNAs. Finally, we implemented alternative RNA-targeting CRISPR-Cas systems with reduced or absent collateral activity. Altogether, these findings contribute to CRISPR-Cas technology optimization for RNA targeting in zebrafish through transient approaches and assist in the progression of in vivo applications.

## Introduction

RfxCas13d is a class 2/type VI CRISPR-Cas RNA endonuclease that together with a guide RNA (gRNA) targets RNA by RNA-RNA hybridization. The CRISPR-RfxCas13d system has been efficiently used to eliminate RNA and therefore possesses extraordinary potential for biotechnology and biomedicine^1–6^. We recently optimized this technology *in vivo* using ribonucleoprotein (RNP) complexes or mRNA-gRNA delivery that allows effective, transient, and cytosolic mRNA knockdown (KD) during vertebrate embryogenesis, including zebrafish, medaka, killifish and mouse, among other vertebrate embryos^7–15^. However, through our continued work we have identified a set of limitations that need to be addressed to further expand the *in vivo* capabilities of the CRISPR-RfxCas13d system.

First, targeting nuclear RNAs or zygotically-expressed genes transcribed after gastrulation is less efficient^7^. A second limitation is that a subset of *in vitro* transcribed gRNAs, but not their chemically synthesized versions, can trigger toxic effects during embryogenesis. Thirdly, as described in mammalian cells^16–19^ and in other CRISPR-Cas systems both *in vivo* and *ex vivo*^20–22^, on-target gRNA activity is variable and can challenge the targeting efficiency^7^. Moreover, CRISPR-RfxCas13d specificity has come under scrutiny lately due to the parallel and recent discoveries of collateral activity in eukaryotic cells^23–29^. Collateral activity is a shared feature of all Cas13 family endonucleases and has been well established in bacteria and *in vitro*^30–32^. It is defined as the cleavage of non-target RNAs that relies upon on-target gRNA recognition and stems from the three-dimensional structure of the two HEPN (Higher Eukaryotes and Prokaryotes Nucleotide-binding) nuclease domains in Cas13 family members. After the activation and a conformational change of Cas13, the HEPN domains are exposed on the outside of the protein and are able to cleave other accessible RNA molecules^33,34^. In eukaryotes, this phenomenon has been mainly observed in *ex vivo* contexts such as mammalian or *Drosophila* cell cultures and/or when gRNAs and RfxCas13d are highly expressed from constitutive promoters^23–29^. RfxCas13d collateral activity has yet to be characterized *in vivo* using transient targeting approaches (the delivery of RNP or mRNA-gRNA formulations) or assessed in the context of embryo development.

Here, we have enhanced RNA-targeting CRISPR-Cas technology *in vivo* through different and compatible approaches using zebrafish embryos as a model system. First, we show that chemically modified gRNAs, along with *RfxCas13d* mRNA, significantly increase the loss-of-function phenotype penetrance when targeting mRNA from genes with late expression during development. Second, we have implemented an approach to select high-quality *in vitro* transcribed gRNAs to avoid potential toxic effects *in vivo*. Third, we optimize RfxCas13d nuclear targeting along zebrafish early development by incorporating nuclear localization signals previously used to increase the activity of DNA-targeting CRISPR-Cas systems. Fourth, we compare different computational models recently developed in mammalian cell cultures^17–19^, analyze their accuracy to predict the activity of 200 gRNAs delivered as RNP complexes and define the most accurate approach for classifying CRISPR-RfxCas13d efficiency *in vivo*. Fifth, we demonstrate that transient CRISPR-RfxCas13d approaches can be used to deplete the vast majority of naturally present mRNAs in zebrafish embryos without inducing collateral activity, although this effect is triggered when targeting extremely abundant and ectopic mRNAs. Finally, we evaluate and compare the on-target and collateral activity of other RNA-targeting CRISPR-Cas systems, such as CRISPR-Cas7-11, CRISPR-DjCas13d and a high-fidelity version of RfxCas13d, formulated as RNP complexes. We demonstrate that CRISPR-Cas7-11 and specially CRISPR-DjCas13d can efficiently eliminate mRNAs *in vivo* and with absent or lower collateral effects than CRISPR-RfxCas13d, respectively when depleting highly abundant and ectopic RNAs in zebrafish embryos. Overall, our work constitutes a significant contribution towards better comprehension and enhancement of transient CRISPR-Cas approaches to target RNA in zebrafish embryos that will ultimately facilitate more effective integration of RNA-targeting CRISPR-Cas into *in vivo* KD-based biotechnological and biomedical applications.

## Results

### CRISPR-RfxCas13d guide RNA optimizations lead to straight-forward and sustained targeting in zebrafish embryos

CRISPR-RfxCas13d has shown high activity when targeting maternally provided and early transcribed mRNAs during vertebrate embryogenesis^7–14,35^. However, the system performed with lower efficiency when depleting genes expressed later during development after 7-8 hours post-fertilization (hpf)^7^. Since chemically modified gRNAs (cm-gRNAs) have been used to maintain the efficiency of mRNA depletion in mammalian cell cultures^36^, we hypothesized that this could be used during zebrafish embryogenesis to sustain and increase RNA targeting *in vivo*. We employed the most efficient chemical modification described in *in vitro* approaches (cm-gRNAs, 2’-O-methyl analogs and 3’-phosphorothioate internucleotide linkage in the last three nucleotides, Mendez-Mancilla et al., 2022^36^) combined with either mRNA or purified protein RfxCas13d (**Fig. 1A**). We targeted different maternally-provided mRNAs and zygotically-transcribed mRNAs in zebrafish embryos whose lack of function and phenotype penetrance could be easily visualized and quantified^7,8,37^ (**Fig. 1B-E, Extended Data Fig. 1**). For example, the KD of *nanog* mRNA leads to epiboly defects with a substantial fraction of embryos at 30-50% epiboly at 6 hpf instead of germ ring or shield stage. The KDs of *tbxta* (hereafter named as *no-tail*) and *noto* mRNAs cause a reduction and/or the lack of the notochord and posterior part in the embryo (**Fig. 1C**). In addition, the KD of *rx3* impairs eye development, and the KDs of *tyrosinase* (*tyr*), *slc45a2* (hereafter named as *albino*) and *slc24a5* (hereafter named as *golden*) mRNAs produce a loss of pigmentation (**Fig. 1C, Extended Data Fig. 1A**). While maternally-provided *nanog* mRNA was efficiently targeted by RNP complexes without any improvement using cm-gRNAs (**Fig. 1D**), early zygotically-expressed RNAs (*no-tail* and *noto*) KD experienced a subtle but still significant increase in the phenotype penetrance (**Fig. 1D, Extended Data Fig. 1B**). Conversely, independently of the use of cm-gRNAs, RNP complexes were much less active when targeting mRNAs from genes whose main expression occurs later during development after 7-8 hpf (**Fig. 1B, Extended Data Fig. 1B**). Notably, the combination of *RfxCas13d* mRNA and cm-gRNA showed a superior efficiency eliminating these mRNAs with a more penetrant phenotype and robust targeting (**Fig. 1E, Extended Data Fig. 1C-D**). Altogether, these results demonstrate that cm-gRNAs increase the activity of CRISPR-RfxCas13d on transcripts from genes expressed later during zebrafish embryogenesis and validate the use of cm-gRNA to deplete RNA *in vivo*. Additionally, we tested whether longer spacers (30 nucleotides, nt) might slightly boost RNA depletion *in vivo* by CRISPR-RfxCas13d as it was previously described in mammalian cell cultures^16^. However, either 23 or 30 nt gRNAs led to a similar efficiency targeting maternal or zygotically expressed mRNAs with both RfxCas13d mRNA or protein (**Extended Data Fig. 2**).

**Figure 1.**
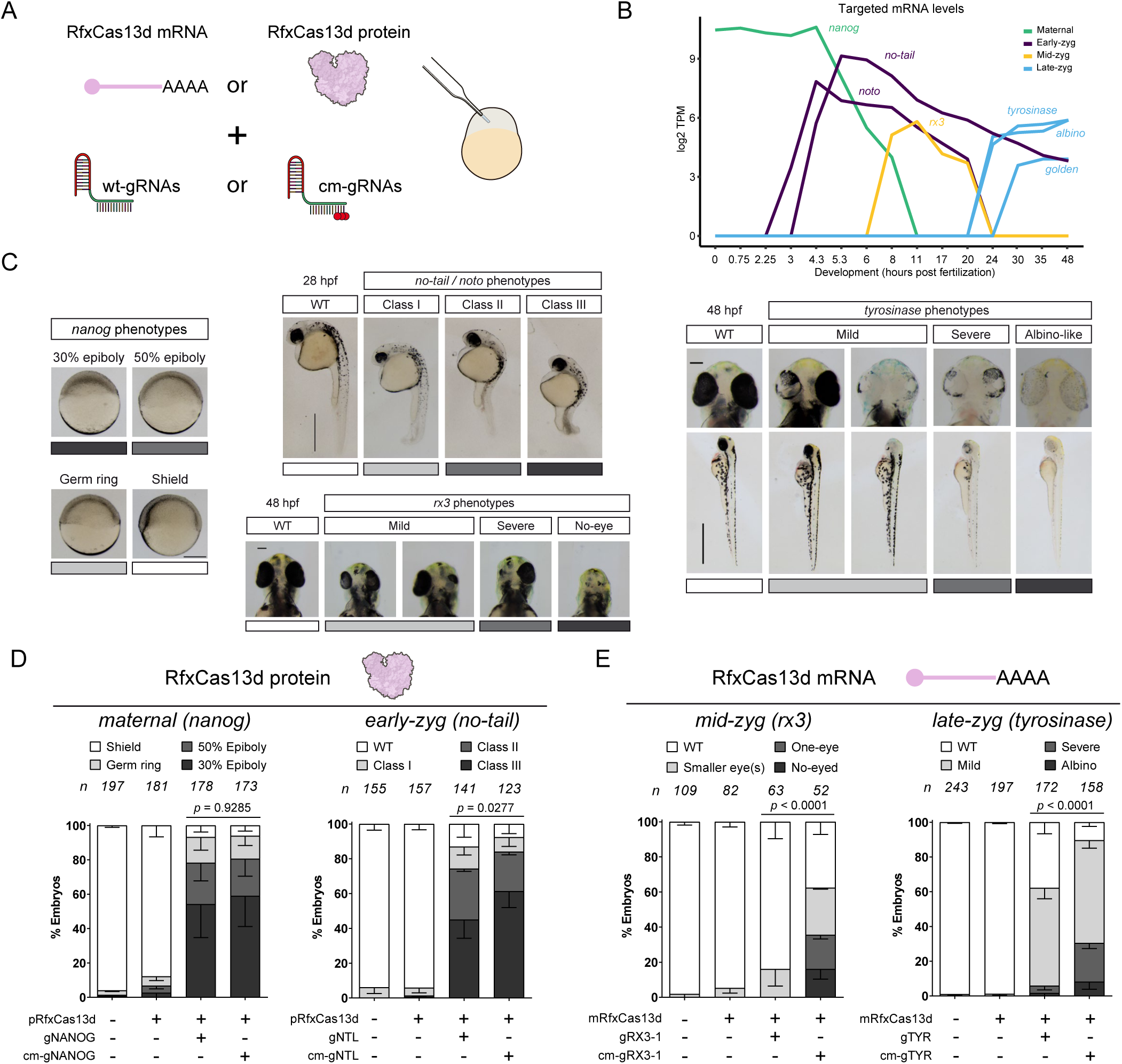
Sustained CRISPR-RfxCas13d targeting in zebrafish embryos. **A)** Schematic illustration of the experimental setup used to compare CRISPR-RfxCas13d activity using standard (wt-gRNAs) or chemically modified (cm-gRNAs) gRNAs, with both mRNA (*RfxCas13d* mRNA) and purified protein (RfxCas13d protein). One-cell stage zebrafish embryos were injected with 300 pg of RfxCas13d mRNA or 3 ng of RfxCas13d protein together with 1 ng of a mix of three gRNAs (∼300 pg from each gRNA) (otherwise indicated in the corresponding figure legend). **B)** Expression levels (log2 TPM: Transcript per million reads) during the first 48 hours post-fertilization (hpf) of the different maternally-provided (*nanog*) and zygotically-expressed (*tbxta* – *no-tail*, *noto*, *rx3*, *tyrosinase*, *slc45a2* – *albino*, and *slc24a5* – *golden*) mRNAs targeted in this study. Early-zyg, Mid-zyg and Late-zyg indicate early, mid and late zygotically-expressed mRNAs, respectively. Data from zebrafish Expression Atlas^37^. TPM values lower than 10 were referred as 0. **C)** Representative images of the phenotypes obtained after the injection of the mRNA-gRNAs or ribonucleoprotein (RNP) complexes targeting the indicated maternally-provided or zygotically-expressed mRNAs. *Nanog* loss-of-function phenotypes were evaluated at ∼6 hpf: 30% epiboly, 50% epiboly, germ ring, and shield stages correspond to 4.6, 5.3, 5.7, and 6 hpf in non-injected embryos growing in standard conditions, respectively (scale bar, 0.25 mm). *Tbxta* – *no-tail* or *noto* loss-of-function phenotypes were evaluated at 28 hpf (scale bar, 0.5 mm). Class I: short tail (least extreme), Class II: absence of notochord and short tail (medium level), and Class III: absence of notochord and extremely short tail (most extreme). *Rx3*, loss-of-function phenotypes were evaluated at 48 hpf (scale bar, 0.1 mm). Mild: smaller eye(s) (least extreme), Severe: absence of one eye (medium level), and No-eye: absence of both eyes (most extreme). *Tyrosinase* loss-of-function phenotypes (reduced or lack of pigmentation) were evaluated at 48 hpf (scale bar, 0.75 mm for lateral views and 0.1 mm for insets). Mild (least extreme), Severe (medium level), and Albino-like (most extreme). **D)** Stacked barplots showing percentage of observed phenotypes in zebrafish embryos injected with standard or cm-gRNAs targeting *nanog* (gNANOG) and *no-tail* (gNTL) together with RfxCas13d protein (pRfxCas13d). (n) total number of embryos is shown for each condition The results are shown as the averages ± standard deviation of the mean of each phenotypic category from at least two independent experiments. Chi-square (χ^2^) statistical tests were performed to compare standard and cm-gRNAs. **E)** Stacked barplots showing percentage of observed phenotypes in zebrafish embryos injected with standard or cm-gRNAs targeting *rx3* (one single gRNA, gRX3-1) and *tyrosinase* (gTYR) together with RfxCas13d mRNA (mRfxCas13d). (n) total number of embryos is shown for each condition. The results are shown as the averages ± standard deviation of the mean of each phenotypic category from at least two independent experiments. χ^2^ statistical tests were performed to compare standard and cm-gRNAs.

In all previous experiments (**Fig. 1 and Extended Data Fig. 1 and 2**), we employed chemically synthesized and commercially available gRNAs (see Methods). Alternatively, efficient and specific gRNAs can be produced through oligo-annealing and fill-in PCR, followed by *in vitro* transcription (hereafter: IVTed gRNAs)^7,8^. Nevertheless, employing IVTed gRNAs we have occasionally observed toxic effects in zebrafish embryos. For example, when targeting *si:dkey-93m18.4* mRNA, a lowly expressed transcript, with three individual IVTed gRNAs, we detected three distinct phenotypes when injected with RfxCas13d protein. While gRNA-1 had no effect during embryogenesis, gRNA-2 showed early embryogenesis delay, and gRNA-3 was lethal with previous developmental defects even by 2 hpf (**Fig. 2A, Extended Data Fig. 3A**). Conversely, targeting *si:dkey-93m18.4* mRNA with any of these three gRNAs showed comparably strong (>90%) KD at 4 hpf (**Fig. 2B**). We next wondered whether this toxic effect stemmed from *in vitro* transcription synthesis or was a direct result of gRNA sequence. To address this, we used chemically synthesized gRNA-1 and gRNA-3 targeting *si:dkey-93m18.4* mRNA. Injection of chemically synthesized gRNA-1 produced no developmental effects, as expected. In contrast to the lethality observed with IVTed gRNA-3, embryos injected with chemically synthesized gRNA-3 exhibited normal development (**Fig. 2A, Extended Data Fig. 3A**). Critically, RfxCas13d with chemically synthesized gRNAs achieved similar *si:dkey-93m18.4* knockdown to their IVTed counterparts (**Fig. 2B**). Taken together, this data demonstrates that a subset of RfxCas13d gRNAs can have toxic effects when produced by *in vitro* transcription.

**Figure 2.**
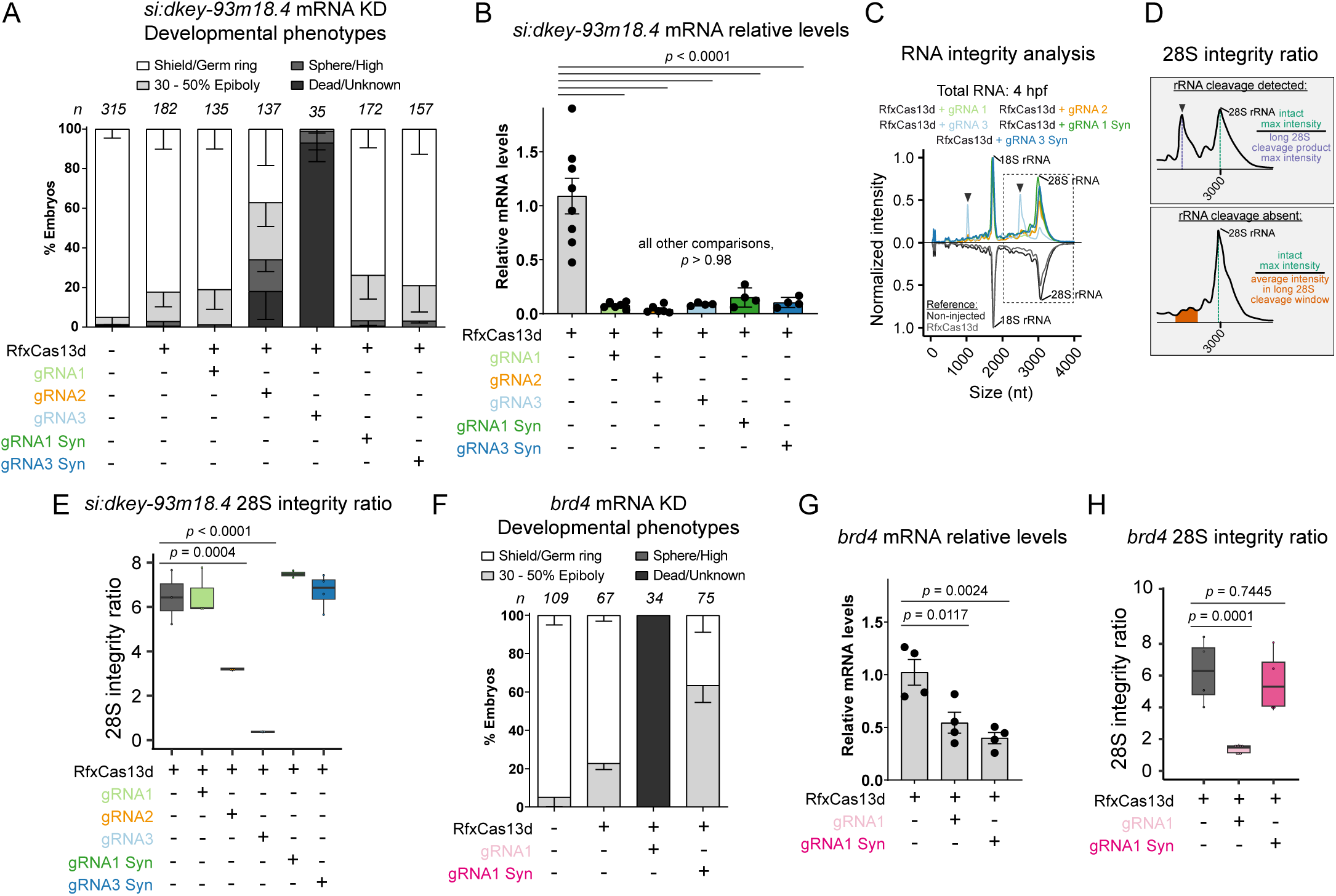
*In vitro* transcribed gRNAs can induce toxicity. **A)** Stacked barplots showing percentage of developmental effect of *si:dkey-93m18.4* KD at 6 hpf using *in vitro* transcribed gRNAs (IVTed, gRNAs 1-3) or chemically synthetized (Syn) gRNAs (400 pg/embryo). (n) total number of embryos is shown for each condition. The results are shown as the averages ± standard deviation of the mean of each developmental stage from at least two independent experiments. **B)** Boxplot showing relative *si:dkey-93m18.4* mRNA levels measured by qRT-PCR at 4 hpf from injected embryos in **A**. Results are shown as the averages ± standard deviation of the mean from at least four biological replicates from two independent experiments. One-way ANOVA followed by Tukey post-hoc analysis was performed. *cdk2ap2* mRNA was used as a normalization control. **C)** Mirrored line-plot represents relative abundance (normalized fluorescence intensity of bioanalyzer electrophoresis) of RNA species present in total RNA at 4 hpf from RfxCas13d mediated *si:dkey-93m18.4* KD and control embryos. Traces are single replicates. Non-injected (gray) and RfxCas13d alone (black) traces are inverted as a reference for RfxCas13d + gRNA1 (light green), RfxCas13d + gRNA2 (orange), RfxCas13d + gRNA3 (light blue), RfxCas13d + gRNA1 Syn (dark green), and RfxCas13d + gRNA3 Syn (dark blue). Black arrows denote peaks from 28S rRNA cleavage. **D)** Cartoon of 28S integrity ratio calculation as the intensity of the 28S rRNA relative to the long 28S cleavage product when was detected (top) and when it was absent (bottom). See Methods for further details. **E)** Boxplot of 28S integrity ratio *in vivo* at 4 hpf from RfxCas13d mediated *si:dkey-93m18.4* KD and control embryos. At least two biological replicates were analyzed. The mean, first and third quartile are represented for each condition. One-way ANOVA followed by Dunnett’s post-hoc analysis was performed. **F)** Stacked barplots showing percentage of developmental effect of *brd4* KD at 6 hpf using an *in vitro* transcribed or chemically synthetized gRNA (IVTed, gRNA1 and Syn, respectively; 300 pg of gRNA/embryo). (n) total number of embryos is shown for each condition. The results are shown as the averages ± standard deviation of the mean of each developmental stage from at least two independent experiments. **G)** Boxplot showing relative *brd4* mRNA levels measured by qRT-PCR at 4 hpf from injected embryos in **F**. The mean, first and third quartile are represented for each condition. Four biological replicates from two independent experiments were analyzed. One-way ANOVA was performed. *taf15* mRNA was used as a normalization control. **H)** Boxplot of 28S integrity ratio *in vivo* at 4 hpf from RfxCas13d mediated *brd4* KD and control embryos. The mean, first and third quartile are represented for each condition. At least four biological replicates were analyzed. One-way ANOVA followed by Dunnett’s post-hoc analysis was performed.

One of the molecular outcomes from these toxic effects was that, beyond the expected 18S and 28S ribosomal RNA (rRNA) observed in an RNA integrity analysis, we detected two prominent RNA species (**Fig. 2C, Extended Data Fig. 3B**) in *si:dkey-93m18.4* KDs consistently of ∼1000 and ∼2500 nucleotides (nt) in length and associated with developmental effects. We employed these two 28S rRNA cleavage species and their positions to ratiometrically quantify the integrity of the 28S rRNA. Controls and *si:dkey-93m18.4* KDs without developmental effects exhibited similar 28S integrity (**Fig. 2D-E**) whereas KDs with IVTed gRNA-2 and gRNA-3 showed significant decreases in 28S integrity that scale with both phenotype onset and severity (**Fig. 2A, C-E, Extended Data Fig. 3A-B**).

Correspondingly, we observed similar results when we targeted *brd4* mRNA using an IVTed or a chemically synthesized gRNA, both inducing a significant reduction of transcript levels (**Fig. 2F-G**). *Brd4* mRNA knockdown causes epiboly defects^7^ that were recapitulated by the chemically synthesized gRNA (**Fig. 2F)**. However, the IVTed gRNA triggered a severe early embryogenesis lethality associated with a 28S rRNA fragmentation (**Fig. 2F and H and Extended Data Fig. 3B).**

Furthermore, we developed a simple *in vitro* rRNA integrity assay method to screen for IVTed gRNAs without a 28S rRNA cleavage effect associated with toxicity. We combined RfxCas13d protein, and/or gRNA with zebrafish total RNA and examined the total RNA through electrophoresis (**Extended Data Fig. 3C**). For example, in *si:dkey-93m18.4* KDs we observed that neither RfxCas13d alone nor gRNA-1 or chemically synthesized gRNAs (1 and 3) as well as gRNAs alone (**Extended Data Fig. 3D**) had any substantial effect on the rRNA integrity in this assay. In contrast, RfxCas13d with IVTed gRNA-2 and gRNA-3 triggered rRNA cleavage *in vitro* (**Extended Data Fig. 3D**), which was more severe than that observed in total RNA from injected embryos (**Extended Data Fig. 3B**).

Altogether, these data demonstrated that some IVTed gRNAs for RfxCas13d may trigger 28S rRNA fragmentation *in vitro* and *in vivo* that is associated with severe defects during embryogenesis and toxicity. Importantly, these toxic effects can be overcome by i) using chemically synthesized gRNAs or ii) pre-screening IVTed gRNAs with our *in vitro* rRNA integrity assay.

### Enhancing nuclear RNA depletion by CRISPR-RfxCas13d in zebrafish embryos

Efficient nuclear RNA targeting can be crucial to eliminate RNAs located in this cellular compartment, such as long non-coding RNAs or primary microRNAs^38–40^. However, our optimized approach triggers mRNA KD in the cytosol. Indeed, a version of RfxCas13d with nuclear localization signals (NLS, RfxCas13d-NLS) was much less active in zebrafish embryos^7^ in contrast to what it was observed in mammalian cell cultures when RfxCas13d targets the nascent mRNA^41^. We hypothesized that the optimization of NLS could improve the efficacy of nuclear RNA elimination mediated by RfxCas13d in zebrafish embryos. We tested 4 NLS formulations that have shown to increase nuclear targeting effectiveness with CRISPR-Cas systems with DNA endonuclease activity^42–45^. All NLS versions were innocuous during early zebrafish development when fused to RfxCas13d (**Extended Data Fig. 4**). We observed that 2 NLS (SV40-Nucleoplasmin long NLS) at the carboxy-terminus^42^ of RfxCas13d (RfxCas13d-2C-NLS) significantly caused the highest phenotype penetrance observed at 48 hpf targeting the primary and nuclear transcript of miR-430 (pri-miR-430) (**Fig. 3A-B**), a microRNA involved in early development regulation that eliminate hundreds of mRNAs during the first hours of development^46–48^. Indeed, this pri-miR-430 targeting induced by RfxCas13d-2C-NLS, specifically triggered a global stabilization of a subset of mRNAs (n=203) whose degradation was more strictly dependent on miR-430^49,50^ (**Fig. 3C**) without any substantial alteration of other maternal mRNA decay programs depending on other maternally-provided or zygotically-expressed factors^50^. Notably, despite the intrinsic mosaicism of the microinjection, pri-miR-430 KD phenotype at 48h hpf was partially rescued by a mature version of miR-430 (**Fig. 3D**), strongly suggesting that observed developmental defects were specifically caused by the miR-430 loss-of-function. In addition, the optimized RfxCas13d-2C-NLS also triggered efficient and significant KD of a small nuclear RNA, *u4atac* snRNA^51^, that was not depleted by cytosolic RfxCas13d or our previous RfxCas13d-NLS^7^ (**Fig. 3E**).

**Figure 3.**
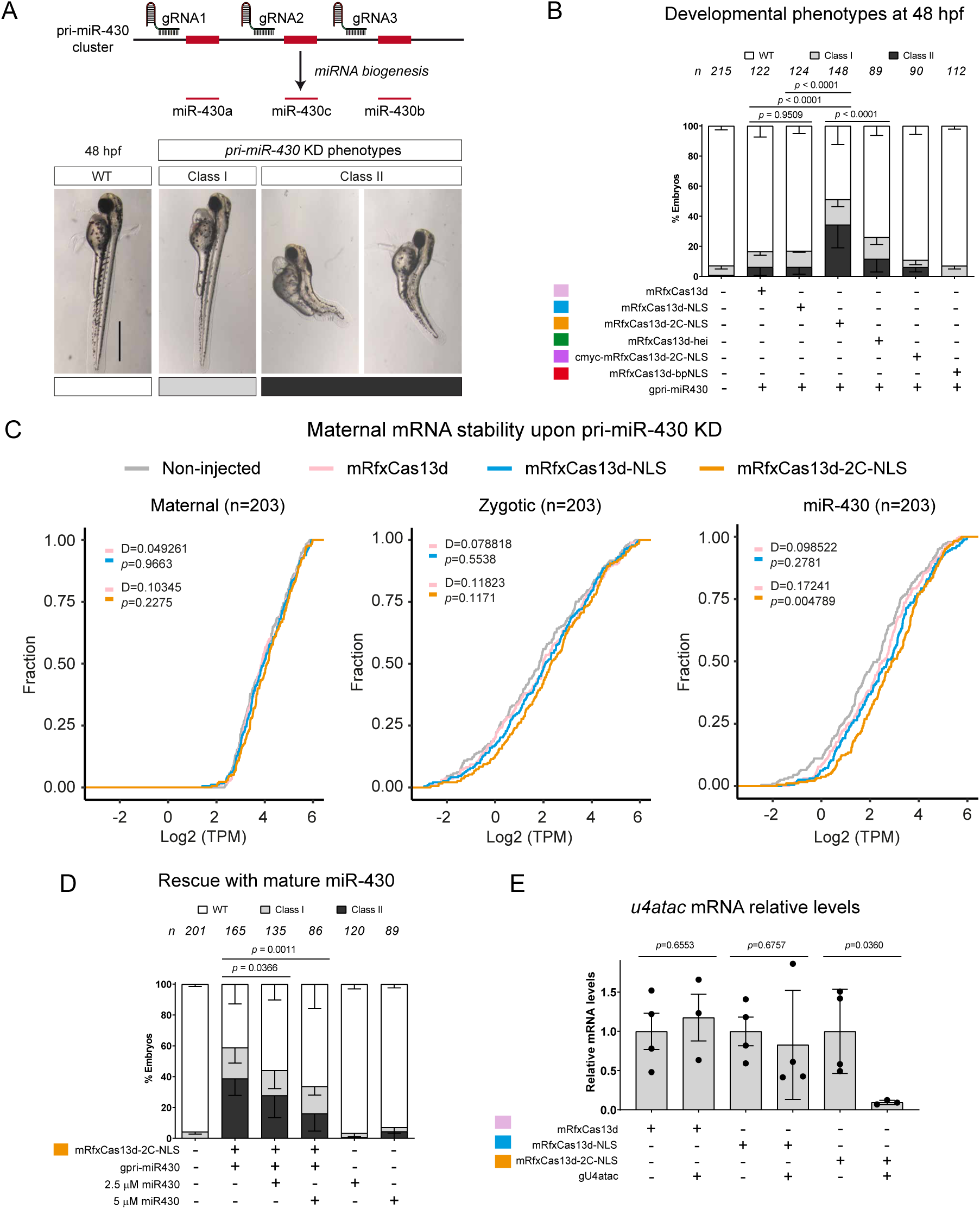
An optimized localization signal enhances CRISPR-RfxCas13d nuclear RNA-targeting. **A)** Diagram illustrating the positions of three gRNAs targeting primary miR430 transcript (pri-miR430), red rectangles indicate the mature microRNAs (miR430a, miR430b, and miR430c). Scheme based on Hadzhiev et al., 2023^48^ (top). Representative images of developmental defects observed at 48 hpf when targeting miR430 (scale bar, 0.75 mm) (bottom). Class I: heart oedema (least extreme), and Class II: body curvature and/or tail blister (most extreme). **B)** Stacked barplots showing percentage of observed phenotypes at 48 hpf comparing cytoplasmic (mRfxCas13d, pink), NLS (mRfxCas13d-NLS, blue), optimized 2C-NLS (mRfxCas13d-2C-NLS, orange), heitag (mRfxCas13d-hei, green), cmyc-2C-NLS (cmyc-mRfxCas13d-2C-NLS, purple) and bpNLS (mRfxCas13d-bpNLS, red) versions of *RfxCas13d* mRNA targeting pri-miR430 (gpri-miR430). Configuration of NLS signals in the different NLS versions are shown in **Extended Data Fig. 4A**. (n) total number of embryos is shown for each condition. The results are shown as the averages ± standard deviation of the mean of each phenotypic category from at least two independent experiments. χ^2^ statistical test was performed to indicated comparisons. **C)** Cumulative plot of the mRNA levels that are degraded by different pathways (maternal, zygotic and miR-430) through early zebrafish development and determined by Vejnar et al., 2019^50^ and Medina-Muñoz et al., 2021^49^ in control (Non-injected; gray line), and different pri-miR430 KD conditions using cytoplasmic (mRfxCas13d; pink line), NLS (mRfxCas13d-NLS; blue line) or optimized NLS (mRfxCas13d-2C-NLS; orange line) versions of RfxCas13d. Number of mRNA controlled by each pathway is shown (n). 203 random mRNAs degraded by maternal and zygotic pathways were selected and *p*-values and distances (D, maximal vertical distance between the compared distribution) were calculated using Kolmogorov-Smirnov tests. **D)** Stacked barplots showing percentage of observed phenotypes at 48 hpf upon pri-miR-430 targeting using mRfxCas13d-2C-NLS with and without a rescue by a mature miR-430. (n) total number of embryos is shown for each condition. The results are shown as the averages ± standard deviation of the mean of each observed phenotype. 1 nL of the indicated concentration of mature miR430 duplexes was injected. χ^2^ statistical tests were performed for indicated comparisons. **E)** Barplots showing snRNA *u4atac* relative levels measured by qRT-PCR at 6 hpf comparing cytoplasmic (mRfxCas13d, pink), NLS (mRfxCas13d-NLS, blue) and optimized NLS (mRfxCas13d-2C-NLS, orange) versions of RfxCas13d. Results are shown as the averages ± standard deviation of the mean from four biological replicates from two independent experiments. T-test statistical analyses were performed to compare control vs KD. miR-430b was used as a normalization control.

Altogether, our results demonstrate that CRISPR-RfxCas13d-2C-NLS system used as a mRNA-gRNA formulation efficiently depletes nuclear RNAs in zebrafish embryos.

### RNAtargeting, an *ex vivo*-based computational model, can moderately predict CRISPR-RfxCas13d activity *in vivo*

Several computational models have been recently developed to predict CRISPR-RfxCas13d activity^16–19^. These models were based on data where RfxCas13d and gRNAs were expressed from constitutive promoters in mammalian cell cultures ^16–19^. However, *in vivo* approaches frequently imply the delivery of a purified RfxCas13d protein or Rfx*Cas13d* mRNA and chemically synthesized or *in vitro* transcribed gRNAs. To examine whether computational models based on cell culture data were able to accurately predict CRISPR-RfxCas13d activity *in vivo*, we measured the efficiency of approximately 200 gRNAs in zebrafish embryos using our optimized and transient approach based on RNP complexes. First, to define the maximal number of gRNAs that could be used in zebrafish embryos allowing a successful detection of highly active gRNAs, we co-injected 10 and 25 gRNAs as optimal and suboptimal targeting conditions (100 pg and 40 pg of gRNA per embryo, respectively) (**Extended Data Fig. 5A-B**). Next, we generated gRNA quintiles (q) based on their activity and observed that up to 25 gRNAs injected together allowed us to detect highly efficient gRNAs (q4 and q5, respectively) previously identified in optimal conditions (**Extended Data Fig. 5**, fold change > 3.5, q4 and q5). Consequently, we injected 8 independent combinations (gRNA set 1 to 8) of 25 gRNAs to KD 75 mRNAs (2-3 gRNAs per transcript) with high-moderate and stable levels between 1 and 4 hpf, where most of the targeting likely occurs (**Fig. 4A, Extended Data Fig. 5C**). Then, we performed a RNA-seq analysis of each gRNA set at 4 hpf (**Extended Data Fig. 6A-B**) and analyzed the efficiency of those gRNAs (n=191) whose activity could be predicted by the most recent and updated computational CRISPR-RfxCas13d models based on the activity of hundreds of thousands constitutively expressed gRNAs in mammalian cell culture^17–19^ (**Fig. 4A** and see Methods for details). Among the assessed models, RNAtargeting^19^ was the most accurate classifying CRISPR-RfxCas13d RNP complexes activity in zebrafish embryos (**Fig. 4B, Extended Data Fig. 6C**). Indeed, RNAtargeting was able to classify 5 out of 8 set of gRNAs with a similar or even more accuracy than what was calculated for an independent *ex vivo* data used as control (Cell culture data, Pearson’s correlation coefficient R> 0.38, **Fig. 4B**). Furthermore, RNAtargeting was the most efficient computational model distinguishing highly (top 5) from poorly (bottom 5) active gRNAs per set (**Fig. 4C-E**). These results suggest that RNAtargeting is the most useful current tool to select competent gRNAs. Nevertheless, RNAtargeting was less accurate at predicting CRISPR-RfxCas13d RNP activity in 3 out 8 sets, especially in one of them where there was not a positive correlation between the predicted scores and *in vivo* activity (**Fig. 4B**; gRNA set 7, Pearson’s correlation coefficient R= −0.05). Together, our results validate the use of RNAtargeting to moderately classify CRISPR-RfxCas13d activity delivered *in vivo* as RNP complexes.

**Figure 4.**
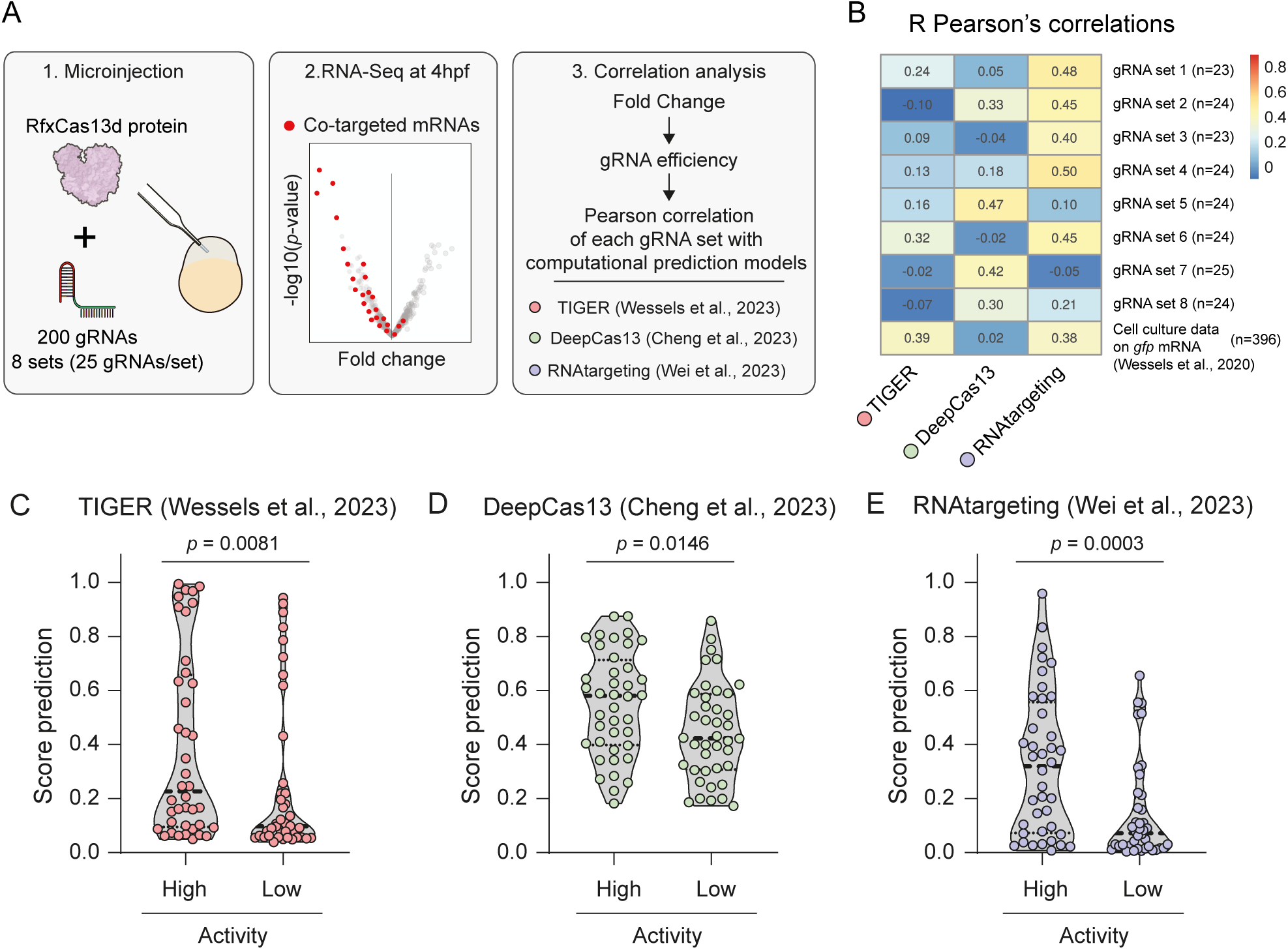
RNA-targeting, an *ex vivo*-based computational model can moderately predict CRISPR-RfxCas13d activity *in vivo*. **A)** Workflow to assess the efficiency of computational models based on *ex vivo* data correlate to predict *in vivo* activity. 1) Diagram illustrating the injection of 8 different sets of 25 gRNAs together with RfxCas13d protein in one-cell stage zebrafish embryos. 2) RNAseq were performed from injected embryos at 4 hpf. Co-targeted mRNAs per set are depicted in red. 3) Fold Changes obtained from RNAseq data were used to calculate each gRNA efficiency and this employed to calculate Pearson correlation with scores obtained from the most recent and updated computational prediction models based on *ex vivo* data (Wessels et al., 2023^17^, Cheng et al., 2023^18^, Wei et al., 2023 Cell Systems^19^). See Methods for further details. **B)** Heatmap of Pearson’s correlations coefficient from each gRNA set with scores predicted by computational models in **A:** TIGER, red dots, Wessels et al., 2023^17^; DeepCas13, green dots, Cheng et al., 2023^18^; RNAtargeting, purple dots, Wei et al., 2023^19^). Pearson’s correlations coefficient per computational model from cell culture data on *gfp* mRNA targeting using 396 gRNAs is included as a reference control (Wessels et al., 2020^16^). **C, D, E)** Violin-plots representing the score of the five highest (High) and five lowest (Low) active gRNAs per set predicted by TIGER^17^ (**C**), DeepCas13^18^ (**D**) and RNAtargeting^19^ (**E**). The median and first and third quartiles of the 40 gRNAs score are represented as dashed and dotted lines, respectively. Non-parametric Mann-Whitney U statistical tests were performed to compare the score of high and low active gRNAs.

### Minimal collateral activity targeting endogenous mRNAs by CRISPR-RfxCas13d

The recently described collateral activity triggered by CRISPR-RfxCas13d induces different molecular outcomes that can be used as hallmarks of this effect. The main consequence of this effect is the uncontrolled elimination of RNAs after the specific and initial targeting of both exogenous and endogenous transcripts^25–29^. Besides, CRISPR-RfxCas13d-induced collateral activity is most severe when targeting highly expressed genes and ultimately causes cell toxicity, a decrease in cell proliferation^25,26,29^ and the cleavage of the 28S rRNA subunit^26,29^. First, and despite that we did not clearly detect any of these effects when previously using our optimized RNP or mRNA-gRNA injections targeting endogenous or ectopic mRNAs^7,8,14^, we sought to investigate the collateral activity when CRISPR-RfxCas13d system targeted a highly concentrated reporter (green fluorescent protein, GFP) mRNA injected in one-cell stage zebrafish embryos. We observed that, for different amounts (from 10 to 100 pg per embryo) of target, CRISPR-RfxCas13d efficiently depleted both mRNA and GFP protein (**Fig. 5A-D**). Notably, embryos injected with more than 50 pg of *gfp* mRNA experienced epiboly defects at 6 hpf (**Fig. 5E**, 30-50% epiboly) when using CRISPR-RfxCas13d as a mRNA-gRNA complex. This effect was more severe when using RNP complexes and, indeed, the KD of 20 or more pg of *gfp* mRNA triggered not only embryogenesis deficiencies but also an arrest during early development and death with the highest concentrations (100 pg per embryo caused a massive embryo death, data not shown) (**Fig. 5F**). Moreover, we observed a cleavage of the 28S rRNA subunit in zebrafish embryos under these conditions correlating with a reduced RNA integrative number (RIN) (**Extended Data Fig. 7A-C**). Interestingly, even with the lowest concentration of *gfp* mRNA (10 pg per embryo), where no developmental delay was noticed upon KD, a notable 28S rRNA fragmentation could be visualized. This result suggests that RNA integrity assay was highly sensitive to detect the collateral activity induced by CRISPR-RfxCas13d (**Extended Data Fig. 7A-B**). Further, when a red fluorescent protein (DsRed) mRNA was co-injected together with CRISPR-RfxCas13d RNP targeting *gfp* mRNA, the fluorescence of both reporters decreased (except for the lowest concentration of *gfp* mRNA), although the developmental and molecular (28S rRNA fragmentation) effects were less severe (**Extended Data Fig. 7D-G**). This could likely be due to a buffer capacity from the *dsred* mRNA that, at high concentration, could partially alleviate the consequences of the collateral activity. To further understand the molecular consequences of the collateral activity during zebrafish embryogenesis, we performed a transcriptome analysis. We observed a global deregulation specifically when targeting an extremely abundant amount of *gfp* mRNA (50 pg/embryo), with 1145 downregulated and 1377 upregulated genes, (**Fig. 5G, Extended Data Fig. 7H)** that correlates with the developmental defects and 28S rRNA fragmentation previously observed. Together, our results demonstrate that RfxCas13d can trigger collateral activity upon the targeting of highly abundant reporter mRNAs.

**Figure 5.**
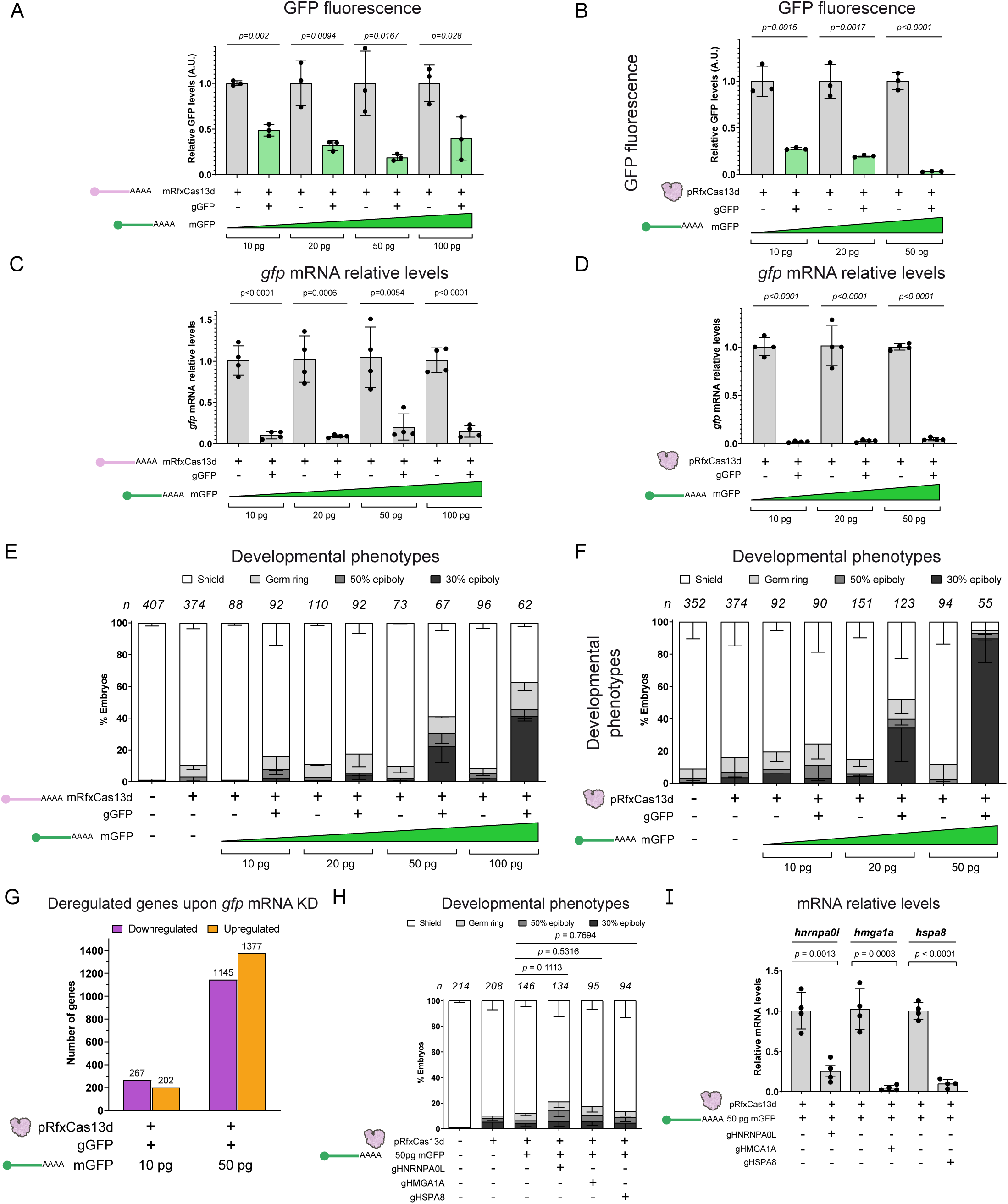
CRISPR-RfxCas13d exhibits physiologically relevant collateral activity only when targeting extremely abundant ectopic mRNAs in zebrafish embryos. **A, B)** Barplots showing GFP fluorescence levels of injected embryos with RfxCas13d mRNA (mRfxCas13d, **A**) or RfxCas13d protein (pRfxCas13d, **B**) together with gRNAs (gGFP) targeting 10, 20, 50 or 100 pg of ectopic *gfp* mRNA (mGFP). Results are shown as the averages ± standard deviation of the mean from three biological replicates of 5 embryos each. T-test statistical analyses were performed, *p*-value is indicated above. **C, D)** Barplots showing *gfp* mRNA relative levels analyzed by qPCR of injected embryos with mRfxCas13d (**C**) or pRfxCas13d (**D**) together with gGFP targeting 10, 20, 50 or 100 pg of ectopic mGFP. Results are shown as the averages ± standard deviation of the mean from four biological replicates from two independent experiments. T-test statistical analyses were performed, *p*-value is indicated above. **E, F)** Stacked barplots representing developmental phenotypes (epiboly stages) of injected embryos with mRfxCas13d (**E**) or pRfxCas13d (**F**) together with gGFP targeting 10, 20, 50 or 100 pg of ectopic mGFP. Representative images of indicated epiboly stages are shown in Fig. 1C. (n) total number of embryos is displayed for each condition. The results are shown as the averages ± standard deviation of the mean of each developmental stage from at least two independent experiments. **G)** Number of down- (purple) and up-regulated (orange) genes from injected embryos with pRfxCas13d together with gGFP and 10 or 50 pg of mGFP. Fold-Change of 2 and *p*-value < 0.001 were set to determine deregulated genes. **H)** Stacked barplots representing developmental phenotypes (epiboly stages) of injected embryos with pRfxCas13d together with 50 pg of mGFP as reporter for collateral activity and gRNAs targeting endogenous *hnrnpa0l*, *hmga1a* and *hspa8* transcripts (gHNRNPA0L, gHMGA1A and gHSPA8, respectively). Representative images of indicated epiboly stages are shown in Fig. 1C. χ^2^ statistical analyses were performed comparing side-by-side each KD condition with embryos injected with pRfxCas13d + mGFP. (n) total number of embryos is displayed for each condition. The results are shown as the averages ± standard deviation of the mean of each developmental stage from at least two independent experiments. **I)** Barplots showing mRNA relative levels analyzed by qRT-PCR at 6 hpf of injected embryos with pRfxCas13d together with 50 pg of mGFP as reporter for collateral activity and gHNRNPA0L, gHMGA1A or gHSPA8. Results are shown as the averages ± standard deviation of the mean from four biological replicates from two independent experiments. T-test statistical analyses were performed comparing side-by-side each KD condition with embryos injected with pRfxCas13d + mGFP. *taf15* mRNA was used as a normalization control.

Next, we sought to analyze whether the collateral activity could be detected when targeting endogenous mRNAs. We selected 3 maternally provided transcripts in zebrafish embryos, all above the top 25 most abundant mRNAs among the polyadenylated mRNAs during the first 6 hpf (**Extended Data Fig. 8A**). When we targeted these mRNAs, we did not observe any developmental defect despite triggering a significant depletion of these endogenous mRNAs (between 75-95% mRNA reduction) (**Fig. 5H-I**). In addition, we did not notice any collateral downregulation of GFP fluorescence from its mRNA that was co-injected together with RNP complexes targeting these endogenous mRNAs (**Extended Data Fig. 8B**). Interestingly, for the depleted endogenous targets, we detected a weak 28S rRNA fragmentation that was much less prominent than observed when targeting the lowest tested amount of *gfp* mRNA (**Extended Data Fig. 7A-B and 8C)**. Nevertheless, this 28S rRNA fragmentation was still significant when rRNA 28S integrity ratio was analyzed (**Fig. 2D and Extended Data Fig. 8D**). This result indicates a minor collateral effect yet without any significant physiological consequence (**Fig. 5H**). Notably, we observed similar results when these endogenous mRNAs were targeted by CRISPR-RfxCas13d RNP complexes without *gfp* mRNA, suggesting that the presence of this transcript did not influence or buffer the collateral effects in these conditions (**Extended Data Fig. 8E-G**). Altogether, our results suggest that the collateral activity from CRISPR-RfxCas13d is minimal and without a physiological relevance even targeting highly abundant mRNAs during zebrafish embryogenesis, but it can be triggered when extremely expressed and ectopic mRNAs such as injected reporters are eliminated.

### Implementation of alternative CRISPR-Cas systems for transient RNA-targeting *in vivo*

Although minimal when targeting endogenous mRNAs, the collateral activity induced by RfxCas13d can still be an issue under certain circumstances *in vivo*. Additionally, the use of RNA-targeting CRISPR-Cas technology as a potential therapeutic application driven by transient approaches needs to ensure a biosafety method where the collateral activity should be minimized or totally absent. Thus, we sought to optimize other RNA-targeting CRISPR-Cas systems *in vivo* that have recently shown a reduced or lack of collateral activity while preserving high on-target efficacy when expressed from constitutive and strong promoters: a high-fidelity version of CRISPR-RfxCas13d (Hf-RfxCas13d^25^), CRISPR-Cas7-11^52^ and CRISPR-DjCas13d^19^. First, we purified these Cas proteins and tested them in zebrafish embryos. None of the protein showed a substantial toxicity when injected alone (**Extended Data Fig. 9A**). Second, we used these Cas proteins to target *no-tail* and *nanog* using 1 and 3 gRNAs per mRNA, respectively. To analyze the efficiency of these systems compared with CRISPR-RfxCas13d, we quantified the phenotype penetrance upon the KD of *nanog* and *no-tail*. While DjCas13d induced a similar phenotype penetrance to RfxCas13d, Hf-RfxCas13d was much less efficient not only when injected as purified protein but also as mRNA (**Fig. 6A-C, Extended Data Fig. 9B-D**), suggesting that this endonuclease, transiently delivered as RNP or mRNA-gRNA complexes, is not highly competent to target mRNA *in vivo*. Further, CRISPR-Cas7-11 recapitulated the expected phenotype from the lack-of-function of *nanog* and *no-tail* with a slightly lower activity than CRISPR-Cas13d systems (**Fig. 6A-C**). Third, we quantified the mRNA levels of *nanog*, *no-tail* and three highly abundant endogenous transcripts (*hnrnpa0l*, *hmga1a*, *hspa8*) formerly analyzed using RfxCas13d (**Fig. 5H-I, Extended Data Fig. 8E-F)** and targeted now by Hf-RfxCas13d, DjCas13d and Cas7-11. Our results validated our previous data based on phenotype analysis **(Fig. 6A-C, Extended Data Fig. 9B-C),** and confirmed that Hf-RfxCas13d showed the lowest efficacy *in vivo* (42.2% average depletion) followed by Cas7-11 (60.1%) and DjCas13d (83.2%) that triggered an efficient mRNA depletion comparable to RfxCas13d activity (80.1%) (**Fig. 6D-I)**. Next, we investigated whether DjCas13d and Cas7-11 showed collateral activity in zebrafish embryos using our previously defined conditions that allow us to measure this effect employing reporter mRNAs. When targeting injected *gfp* mRNA, DjCas13d displayed less severe developmental defects than RfxCas13d. This toxicity increased with the concentration of the target, in agreement with what was observed previously for RfxCas13d in *ex vivo* and *in vivo* conditions^25,26^ (**Fig. 6J-K, Extended Data Fig. 9E, G, I)**. Interestingly, and despite the developmental defects, the fragmentation of the 28S rRNA was not observed (**Extended Data Fig. 9K**) but we detected a non-specific downregulation of DsRed fluorescence when its mRNA was co-injected with high amounts of *gfp* mRNA (**Extended Data Fig. 10A-C**). Notably, and as observed for RfxCas13d, the developmental phenotype triggered by *gfp* mRNA KD was mitigated when *dsred* mRNA was co-injected, suggesting a buffer effect that alleviates this alteration during embryogenesis (**Fig. 6J, Extended Data Fig. 10A-C**). In contrast, Cas7-11 did not show neither developmental defects nor 28S rRNA cleavage when targeting *gfp* mRNA (**Fig. 6J-K, Extended Data Fig. 9F, H, J, L**). We also measured DsRed fluorescence when its mRNA was co-injected together with Cas7-11 RNP and *gfp* mRNA and we did not detect any significant decrease in protein activity (**Extended Data Fig. 10D-F**).

**Figure 6.**
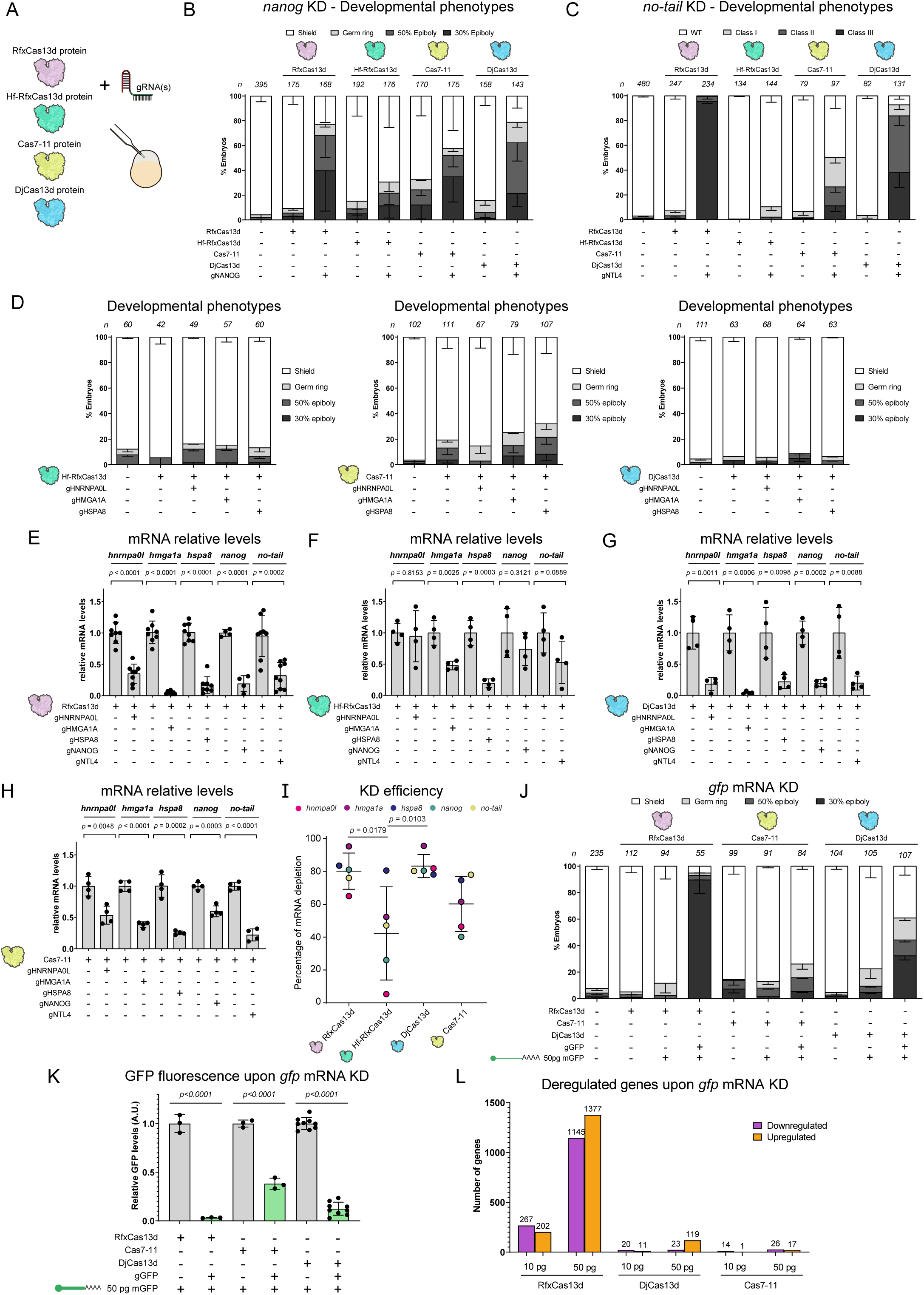
Implementation of alternative CRISPR-Cas systems for transient RNA-targeting *in vivo*. **A)** Schematic illustration of the experimental setup used to compare alternative and high-fidelity CRISPR-Cas RNA targeting systems (Hf-RfxCas13d, DjCas13d and Cas7-11) with CRISPR-RfxCas13d. One-cell stage zebrafish embryos were injected with 3 ng of each purified protein were injected together with 1 ng of a mix of three gRNAs (∼300 from each gRNA) **B, C)** Stacked barplots representing developmental phenotypes from injected embryos with RfxCas13d, Hf-RfxCas13d, Cas7-11 or DjCas13d together with three gRNAs targeting *nanog* (gNANOG, **B**) or one gRNA targeting *no-tail* (gNTL4, **C**). Representative images of epiboly stages and *no-tail* phenotypes are shown in Fig. 1C. (n) total number of embryos is shown for each condition. The results are shown as the averages ± standard deviation of the mean of each developmental stage from at least two independent experiments. **D)** Stacked barplots representing developmental phenotypes (epiboly stages) of injected embryos with Hf-RfxCas13d, Cas7-11 or DjCas13d together with gRNAs targeting endogenous *hnrnpa0l*, *hmga1a* and *hspa8* transcripts (gHNRNPA0L, gHMGA1A and gHSPA8, respectively). Representative images of epiboly stages are shown in Fig. 1C. (n) total number of embryos is displayed for each condition. The results are shown as the averages ± standard deviation of the mean of each developmental stage from at least two independent experiments. **E, F, G, H)** Barplots showing mRNA relative levels analyzed by qRT-PCR at 4 or 6 hpf of injected embryos with RfxCas13d (**E**), Hf-RfxCas13d (**F**), DjCas13d (**G**) or Cas7-11 (**H**) together with gHNRNPA0L, gHMGA1A, gHSPA8, gNANOG or gNTL4. Results are shown as the averages ± standard deviation of the mean from at least four biological replicates from two or more independent experiments. T-test statistical analyses were performed comparing side-by-side each KD condition with embryos injected only with RfxCas13d, Hf-RfxCas13d, DjCas13d or Cas7-11. *hnrnpa0l*, *hmga1a* and *hspa8* relative mRNA levels displayed in Fig. 6E correspond to Fig. 5I and **Extended Data Fig. 8F** data. *taf15* mRNA was used as a normalization control. **I)** Dot-blot showing KD efficiency of RfxCas13d, Hf-RfxCas13d, Cas7-11 and DjCas13d. Each dot represents the mean of the percentage of mRNA depletion analyzed by qRT-PCR from **E-H** panels for the indicated mRNAs in colors; mean and standard deviation are shown. One-way ANOVA followed by Tukey post-hoc analysis was performed, showing *p*-values only for comparisons with significant differences. **J)** Stacked barplots representing developmental phenotypes (epiboly stages) of injected embryos with RfxCas13d, Cas7-11 or DjCas13d together with gRNAs targeting *gfp* (gGFP) and 50 pg of ectopic *gfp* mRNA (mGFP). Representative images of epiboly stages are shown in Fig. 1C. (n) total number of embryos is displayed for each condition. The results are shown as the averages ± standard deviation of the mean of each developmental stage from at least two independent experiments. **K)** Barplots showing GFP fluorescence relative levels at 6 hpf of injected embryos with RfxCas13d, Cas7-11 or DjCas13d together with gGFP and 50 pg of ectopic mGFP. Results are shown as the averages ± standard deviation of the mean from at least three biological replicates of 5 embryos from two independent experiments. T-test statistical analyses were performed, *p*-value is indicated above. **L)** Number of down- (purple) and up-regulated (orange) genes analyzed by RNAseq in injected embryos with RfxCas13d, DjCas13d or Cas7-11 together with gGFP and 10 or 50 pg of mGFP. Fold-Change of 2 and *p*-value < 0.001 were used to determine deregulated genes. Deregulated genes shown for RfxCas13d are from Fig. 5G.

In addition, we performed a whole transcriptomic analysis and observed that while the *gfp* mRNA depletion was highly efficient using either RfxCas13d or DjCas13d (62.68 and 145.01 *gfp* mRNA fold-change, respectively), the transcriptomic deregulation was strongly reduced when employing DjCas13d (a 98% and 91.4% reduction in number of downregulated and upregulated mRNAs in the most extreme condition: 50 pg *gfp* mRNA, **Fig. 6L, Extended Data Fig. 11**). Conversely, *gfp* mRNA depletion mediated by Cas7-11 did not trigger any significant transcriptomic alteration (**Fig. 6L**), but as previously shown (**Fig. 6E-I**), the efficiency of this endonuclease was lower than DjCas13d or RfxCas13d (Cas7-11: 4.26 *gfp* mRNA fold-change, **Extended Data Fig. 11**). Together, our results i) reaffirm that CRISPR-RfxCas13d is an efficient and robust approach to target endogenous mRNA in zebrafish embryos and ii) demonstrate that CRISPR-Cas7-11 and CRISPR-DjCas13d RNP can be used as alternative systems exhibiting a slightly lower or similar on-target activity than CRISPR-RfxCas13d and showing a total absence or reduced collateral effects when targeting extremely abundant ectopic RNAs, respectively.

## Discussion

Our optimized CRISPR-RfxCas13d system has shown a high level of efficiency and specificity targeting maternal and early-zygotically transcribed genes in zebrafish and other vertebrate embryos^7–14^. Here, we have further enhanced CRISPR-Cas RNA-targeting *in vivo* using RNP and mRNA-gRNA formulations in zebrafish as a vertebrate model through different and complementary approaches. First, we have found that chemically modified gRNAs (cm-gRNAs) are able to maintain and increase the activity of CRISPR-RfxCas13d as described in human cells^36^ specially modulating the expression of genes zygotically-transcribed later during development where RNP complexes were less efficient. Although DNA-targeting CRISPR-Cas systems can generate null alleles, this can obscure gene activities that can be compensated through transcriptional adaptation or only detected with a limited gene expression reduction^53–55^. RfxCas13d together with cm-gRNAs allow to titrate mRNA levels of the target that can be an alternative for these scenarios. Indeed, our optimized CRISPR-RfxCas13d system using cm-gRNAs could contribute to widen the applications of combining transient perturbations with single-cell transcriptomics. This approach could allow to simultaneously associate the level of target depletion with a transcriptomic output at single cell level extending the use of this technology recently integrated with zebrafish crispants or F0 mutant embryos targeted by DNA-targeting CRISPR-Cas approaches^56^. Notably, cm-gRNAs have also been recently shown to be useful in zebrafish RNA imaging, expanding their applications *in vivo*^57^.

Second, we have increased nuclear RNA targeting throughout zebrafish embryogenesis using an enhanced NLS formulation. Indeed, we efficiently eliminated nuclear non-coding RNAs such as pri-miR-430 or a snRNA (*u4atac*). The incomplete maturity of the nuclear pore complexes during early zebrafish development that impair nuclear protein import^58^ can affect nuclear DNA or RNA targeting and our results suggest that this may be partially circumvented with an optimized NLS. Notably, this NLS was previously applied to enhance DNA targeting through Cas12a, not only in zebrafish embryos but also in mammalian cells^42^, suggesting that it could be used to improve the nuclear RNA targeting of other CRISPR-Cas systems in different *in vivo* and *ex vivo* models.

Third, we have demonstrated that CRISPR-RfxCas13d RNP activity *in vivo* can be classified, and highly active gRNAs can be predicted by RNAtargeting, a computational model based on mammalian cell culture data, yet with a lower accuracy that shown for approaches with a constitutive expression of both RfxCas13d and gRNA^19^ (**Fig 4B**). However, RNAtargeting is based on the activity from 127000 gRNAs whose spacers were 30 nt long^19^. Although we have shown that RNP activity *in vivo* is similar employing gRNAs with either short (23 nt) or long (30 nt) spacers (**Extended Data Fig. 2**) and we have precisely adapted RNAtargeting to predict our data set based on short spacers (**Fig 4B and see Methods for details**), we can not rule out that the prediction of CRISPR-RfxCas13d RNP activity *in vivo* could be improved when using long spacers. Nevertheless, it has been previously reported that computational models predicting CRISPR-Cas DNA-targeting activity notably differ depending on the employed method. Thus, CRISPR-SpCas9 efficiency prediction models depend on whether the gRNA is constitutively transcribed from a U6 promoter or generated *in vitro*^59^. Considering this and previous CRISPR-Cas activity analysis^20,59^, we speculate that a computational model based on RNP complexes or mRNA-gRNA formulations could outperform the predictive power of RNAtargeting for CRISPR-RfxCas13d activity in zebrafish embryos and enhance the accuracy when selecting highly active gRNAs *in vivo*. Beyond zebrafish embryos, this model could be additionally applied for other biotechnological or biomedical applications where these transient approaches could be used.

Fourth, we have characterized the collateral activity of CRISPR-RfxCas13d in zebrafish embryos previously detected *in vitro* and *in vivo*^23,25–29^. We have now analyzed this effect in different molecular and physiological contexts by employing transient RNA-targeting approaches such as RNP complexes and RfxCas13d mRNA-gRNA formulations during early zebrafish embryogenesis. Importantly, we confirmed that CRISPR-RfxCas13d is very specific, and relevant collateral effects only occur when targeting extremely abundant ectopic RNAs such as *gfp* mRNA. We found that the toxicity and the physiological consequences of the collateral activity increased with the concentration of the *gfp* transcript as described in mammalian cells mRNA^25,26^ (*i.e.* 50 pg of *gfp* mRNA injected per embryo: 10000 TPM at 6 hpf, **Fig. 5E-G**). Interestingly, despite the depletion of lower amounts of *gfp* mRNA (*i.e.* 10 pg per embryo) caused a deregulation of the transcriptome and 28S rRNA fragmentation, it did not induce a developmental defect. Indeed, targeting highly abundant endogenous mRNAs triggered an even fainter 28S rRNA fragmentation and did not generate an early embryogenesis alteration either (**Fig. 5H-I**, **Extended Data Fig. 8C-D**). These findings suggest that ectopic RNAs can elicit much more collateral and toxic effects *in vivo* than endogenous transcripts as reported in mammalian cell cultures^25,28^. Nevertheless, when extremely abundant mRNAs are targeted by CRISPR-RfxCas13d RNP complexes in zebrafish embryos, a simple *in vitro* 28S rRNA cleavage assay can be used to detect even a weak collateral activity without developmental defects (**Extended Data Fig. 8C-D**). In addition, a subset of *in vitro* transcribed gRNAs can trigger toxicity *in vivo* that resembles collateral activity consequences observed when targeting extremely abundant and ectopic mRNA. Whether this effect is certainly CRISPR-RfxCas13d collateral activity remains to be clarified. However, this deleterious response is specifically due to the *in vitro* transcription reaction since the same gRNA when chemically synthesized did not generate any toxic effect while maintaining a similar efficiency of mRNA depletion (**Fig. 2, Extended Data Fig. 3**). In contrast, chemically synthesized are more expensive than *in vitro* transcribed gRNAs and their use can be limited due to the higher cost. Therefore, in case of using *in vitro* transcribed gRNAs we have optimized a fast and straight-forward *in vitro* assay to determine their potential toxic activity that allows to screen for reliable gRNAs to be used *in vivo*. Why this subset of gRNAs induces toxicity during zebrafish embryogenesis depending on whether they are chemically synthesized or *in vitro* transcribed remains to be determined.

Finally, and as an *in vivo* alternative to RfxCas13d specially when targeting extremely abundant transcripts, we have compared the activity of three RNA-targeting CRISPR-Cas systems that showed less or absent collateral activity when gRNAs and Cas were expressed from strong and constitutive promoters^19,25,52^. While Hf-RfxCas13d^25^ exhibited low targeting activity (**Fig. 6A-C and I, Extended Data Fig. 9B-D**), Cas7-11^52^ and DjCas13d^19^ were more active, the latter showing an efficiency comparable to RfxCas13d (**Fig. 6A-C and I**). Interestingly, a recent preprint pointed out that Hf-RfxCas13d activity was inferior than initially described likely due to the lower level of expression of this endonuclease used in this report^60^. Further, this preprint revealed that the concentration of CRISPR-RfxCas13d reagents in mammalian cell culture is crucial to induce collateral activity^60^. Strikingly, we recapitulated these results using a limited amount of CRISPR-RfxCas13d RNP or mRNA-gRNA complexes that show an absence of toxic effects *in vivo* when targeting endogenous mRNAs^14^ (**Fig. 5H and I, Extended Data 8**). Nevertheless, DjCas13d and Cas7-11 reduced or abolish, respectively, the developmental defects during zebrafish embryogenesis and the transcriptomic deregulation observed when RfxCas13d was used to eliminate ectopic and highly expressed RNAs (**Fig. 6J-L, Extended Data Fig. 9E-L and Extended Data Fig. 11**). However, DjCas13d did not totally lack the collateral activity when targeting extremely abundant and ectopic mRNAs inducing a delay during early zebrafish embryogenesis and a decrease in fluorescence levels from a co-injected *DsRed* mRNA control reporter. Interestingly, DjCas13d did not trigger a detectable 28S rRNA cleavage in these conditions, suggesting that this uncontrolled effect may be more efficiently or specifically generated by RfxCas13d. An additional alternative to CRISPR-Cas13 is type III CRISPR-Cas system based on Csm complexes that have been recently employed to target RNA in human cells with minimal off-targets^61^. Although similar approaches have been used in zebrafish embryos^62^, the multicomponent factor of this CRISPR-Cas system with several proteins forming a functional complex challenges its use as RNP particle or mRNA-gRNA formulation *in vivo*. Another possibility to avoid collateral activity may be the application of catalytically dead versions of Cas13 together with translational inhibitors to silence mRNA expression instead of degrading them^63,64^. Instead, we have focused on the characterization, optimization and comparison of recently described high-fidelity CRISPR-Cas systems to eliminate RNA but in an *in vivo* context and using transient formulations that could be employed in potential RNA-editing therapies^65–72^. Further, it has been described that RfxCas13d induces an immune response in humans^73^ that could challenge the biomedical applications of this technology. Whether DjCas13d or Cas7-11 trigger lower immunological effects is something that remains to be determined. In summary, we demonstrate that CRISPR-RfxCas13d is a robust, specific and efficient system to target RNAs in zebrafish embryos but both CRISPR-Cas7-11 and CRISPR-DjCas13d RNP complexes could also be employed specially when targeting exceptionally abundant RNAs, being Cas7-11 less efficient than DjCas13d, although the latter can still trigger developmental defects but in a lower degree than RfxCas13d.

Altogether, our optimizations will not only contribute to broaden and enhance the use of RNA-targeting CRISPR-Cas approaches in zebrafish but also will pave the way to optimize this technology *in vivo* for multiple biomedical and biotechnological applications.

## Methods

### Zebrafish maintenance

All experiments performed with zebrafish conform to national and European Community standards for the use of animals in experimentation and were approved by the Ethical committees from the Pablo de Olavide University, CSIC and the Andalucian Government. Zebrafish wild type strains AB/Tübingen (AB/Tu) or *Tupfel long fin* (TLF) were maintained and bred under standard conditions^74^. Natural mating of wild-type AB/Tu/TLF zebrafish adults (from 6 to 18 months) was used to collect the embryos for subsequent experiments. Selection of mating pairs was random from a pool of 10 males and 10 females. Zebrafish embryos were staged in hours post-fertilization (hpf) as described by Kimmel *et al*. (1995)^75^.

### Guide RNA design and mRNA generation

To design guide RNAs (gRNAs) used in this study (**Extended Data Table 1**), target mRNA sequences were analyzed *in silico* using RNAfold software^76^ (http://rna.tbi.univie.ac.at//cgi-bin/RNAWebSuite/RNAfold.cgi) to select protospacers of 23 (or 30) nucleotides with high accessibility (low base-pairing probability from minimum free energy predictions) to generate gRNAs. All designed gRNAs were synthesized by Synthego (Synthego Corp., CA, USA). Three gRNAs targeting the same mRNA were co-injected, otherwise it is specified in figure legends. *In vitro* transcribed (IVTed) RfxCas13d gRNAs were generated like previously described^7,8^ but the amount of primers were reduced by ten-fold in the fill in PCR (5 μL of 10 μM universal oligo and 5 μL of 10 μM of specific oligos for a 50 μL fill-in PCR).

Heitag (RfxCas13d-hei) version^45^ of RfxCas13d was generated by PCR with Q5 High-Fidelity DNA polymerase (M0491, New England Biolabs) and primers hei-tag_13d_NcoI_fwd and hei-tag_13d_SacII_rev containing a cmyc tag, an oNLS and NcoI site in forward primer and an oNLS and SacII site in reverse primer, and cloned into pT3TS-MCS (Addgene plasmid #31830) backbone after digestion with restriction enzymes NcoI and SacII and ligation with T4 DNA ligase (M0202, New England Biolabs). Similarly, bpNLS (mRfxCas13d-bpNLS; Liang et al., 2022^27^), optimized 2C-NLS (mRfxCas13d-2C-NLS; Liu et al., 2019^42^) and cmyc-2C-NLS (cmyc-mRfxCas13d-2C-NLS; Wu et al., 2019^43^) versions of *RfxCas13d* mRNA were created by PCR using primers NcoI_13d_bpNLS_fwd / 13d_bpNLS_SacII_rev, NcoI_13d_fwd / 13d_2C-NLS_SacII_rev and NcoI_13d_cmyc_fwd / 13d_2C-NLS_SacII_rev to add bipartite NLS signals in N- and C-terminal ends, SV40 and nucleoplasmin long (NLP) NLS signals in C-terminal end, and cmyc tag in N-terminal end and SV40 and NLP NLS signals in C-terminal end, respectively. Each fragment was then cloned into pT3TS-NLS-RfxCas13d (Addgene plasmid #141321) backbone after digestion with restriction enzymes NcoI and SacII. All primers are listed in **Extended Data Table 2**. High-fidelity version of RfxCas13d (Hf-RfxCas13d) was generated by site-directed mutagenesis using QuikChange Multi Site-Directed Mutagenesis kit (Agilent), following manufacturer’s instructions, and replicating the amino acid changes described in Tong et al. (2023^25^; N2V8 version). Primers are listed in **Extended Data Table 2**.

To generate *RfxCas13d*, *RfxCas13d-NLS*, *RfxCas13d-hei*, *RfxCas13d-bpNLS*, *RfxCas13d-2C-NLS*, *cmyc-RfxCas13d-2C-NLS* and *Hf-RfxCas13d* mRNAs, the DNA templates were linearized using XbaI and mRNA was synthesized using the mMachine T3 kit (Ambion) for 2 h. *In vitro* transcribed mRNAs were then DNAse-treated for 20 min with TURBO-DNAse at 37 °C, purified using the RNeasy Mini Kit (Qiagen) and quantified using Nanodrop^TM^ 2000 (Thermo Fisher Scientific).

### Protein purification

The expression vector pET28b was used to clone genes for Hf-RfxCas13d, DjCas13d and Cas7-11 proteins and were transformed into *E. coli* Rosetta DE3 pRare competent cells (70954, EMD Millipore). DjCas13d ORF was codon optimized for zebrafish using iCodon^77^ (www.iCodon.org), purchased from IDT (https://eu.idtdna.com/) and cloned into pET28b vector after NcoI / NotI restriction enzyme digestion. Cas7-11 ORF was PCR amplified from pDF0229- DiCas7-11 (Addgene plasmid #172506) with Q5 High-Fidelity DNA polymerase (M0491, New England Biolabs; primers listed in **Extended Data Table 2**) and cloned into pET28b. A freshly transformed colony was picked and grown over night at 37 °C in LB supplemented with kanamycin and chloramphenicol. This culture was diluted 100 times and grown at 37 °C until an OD_600_ = 0.5 was reached. At this point, cultures for Hf-RfxCas13d and DjCas13d expression were induced with IPTG at a final concentration of 0.1 mM and incubated 3 h at 37 °C. Culture for Cas7-11 expression was cooled for 30 minutes at 4 °C, induced with IPTG at 0.3 mM and cultured over night at 18 °C. Protein purification was performed as described in Hernández-Huertas *et al.* (2022^8^).

### Zebrafish embryo microinjection and image acquisition

One-cell stage zebrafish embryos were injected with 1-2 nL containing 300-600 pg of Cas13(s) mRNA or 3 ng of purified Cas protein and 300-1000 pg of gRNA (see figure legends for details in each experiment). From 10 to 100 pg of ectopic *gfp* mRNA and 75 pg of ectopic *dsRed* mRNA were injected for collateral activity determination. Cas proteins and gRNAs were injected in two rounds to maximize the amount of protein and gRNA (at the indicated concentrations) per injection. Sequences of used gRNAs are indicated in **Extended Data Table 1.**

For the miR-430 rescue experiment, mature miR-430 duplexes (**Extended Data Table 2**) were purchased from IDT (https://eu.idtdna.com/) and resuspended in RNAse-free water. Embryos were injected with 1 nL of 2.5 or 5 µM miR-430 duplex solution (equimolar mix of 3 duplexes miR-430a, miR-430b and miR-430c), using single-use aliquots.

Zebrafish embryo phenotypes and fluorescent pictures were analysed using an Olympus SZX16 stereoscope and photographed with a Nikon DS-F13 digital camera. Images were processed with NIS-Elements D 4.60.00 software. Phenotypes were quantified at 6, 24 or 48 hours post fertilization. GFP and DsRed fluorescence were quantified using Fiji (Image J) software, using 15 embryos separated in three different images (5 embryos per quantified image). Images were converted to 16-bit, background fluorescence was subtracted and fluorescence levels were referred to the ones from embryos injected only with reporter mRNAs.

### Protein sample preparation and Western Blot

Twenty embryos were collected at 6 hours post injection and washed twice with deyolking buffer (55 mM NaCl, 1.8 mM KCl, and 1.25 mM NaHCO_3_). Then, samples were incubated for 5 min with orbital shaking and centrifuged at 300 g for 30 s. Supernatant was removed and embryos were washed with buffer (110 mM NaCl, 3.5 mM KCl, 10 mM Tris-HCl pH 7.4, and 2.7 mM CaCl_2_). Finally, embryos were centrifuged again, and the supernatant was removed. The pellet was resuspended in SDS-PAGE sample buffer (160 mM Tris-HCl pH 8, 20% Glycerol, 2% SDS, 0.1% bromophenol blue, 200 mM DTT).

Sample separation by SDS-PAGE electrophoresis was performed using 10% TGX Stain-Free^TM^ Fast Cast^TM^ Acrylamide Solutions (Bio-Rad). After electrophoresis, protein gel was activated in a Chemidoc MP (Bio-Rad) and blotted onto a nitrocellulose membrane using the Trans-Blot Turbo Transfer System (Bio Rad). The membrane was blocked for 1 h at room temperature in Blocking Solution (5% fat free milk in 50 mM Tris-Cl, pH 7.5, 150 mM NaCl (TBS) with 1% Tween20). Primary antibody Anti-HA (11867423001, Roche) and secondary antibody anti-mouse HRP-labelled (A5278, Sigma-Aldrich) were diluted 1:1000 and 1:5000 respectively in Blocking Solution. The membrane was incubated in primary antibody solution overnight at 4 °C. After primary antibody incubation, the membrane was washed three times in TBS with 1% Tween 20 (TTBS) for 10 min and incubated with the secondary antibody for 60 min at room temperature. Washes were performed as with primary antibody. The protein detection was done with Clarity^TM^ Western ECL Substrate (Bio-Rad) and images were acquired using a ChemiDoc MP (Bio-Rad).

### qRT-PCR

Ten zebrafish embryos per biological replicate were collected and snap frozen in liquid nitrogen to analyze the expression level of the targeted mRNAs by qRT-PCRs at the described hours post injection in figures or legends. Total RNA was isolated using PRImeZOL^TM^ Reagent protocol as described in the manufacturer’s instructions (Canvax Biotech). The cDNA was synthesized from 1000 ng of total RNA using iScript cDNA synthesis kit (Bio-Rad), following the manufacturer’s protocol. cDNA was 1/5 diluted and 2 µl was used per sample in a 10 µl reaction containing 1.5 µl of forward and reverse primers (2 mM each; **Extended Data Table 2**), 5 µl of SYBR® Premix Ex Taq (Tli RNase H Plus) (Takara) and run in a CFX connect instrument (Bio-Rad). PCR cycling profile consisted in a denaturing step at 95 °C for 30 s and 40 cycles at 95 °C for 10 s and 60 °C for 30 s. *taf15* or *ef1α* mRNAs stably expressed along early development were used as controls. To quantify *si:dkey-93m18.4* mRNA levels, 15 embryos per replicate were collected and snap-frozen in tubes containing 350 μL of TRIzol. Total RNA isolation was performed with the Zymo Direct-zol Microprep kit according to manufacturers recommendations (including the DNase digestion step), eluting in 20 μL of nuclease free H2O. Superscript IV was used for reverse transcription of ∼1 μg of total RNA, and resulting cDNA was diluted 1:20 for RT-qPCR. RT-qPCR was run in technical triplicate on a 384-well plate setup by a Tecan robot and run on a QuantStudio 7 workstation usingPerfeCTa® SYBR® Green FastMix® (Quantabio). *cdk2ap2* mRNA stably expressed along early development was used as control. Gene specific oligos for *si:dkey-93m18.4* and *cdk2ap2* are listed in **Extended Data Table 2**.

To analyze the relative levels of *u4atac* snRNA, 15 embryos per replicate were snap frozen and RNA samples enriched in small RNAs were obtained with *mir*Vana mRNA Isolation Kit (AM1561, ThermoFisher Scientific) following the manufacturer’s protocol. cDNA was synthesized from 100 ng of RNA using iScript Select cDNA synthesis kit (Bio-Rad), following the manufacturer’s protocol and using the following miRNA universal primer (5’- GCAGGTCCAGTTTTTTTTTTTTTTTCTACCCC-3’), 2 µl of cDNA were used per sample in a 10 µl reaction containing 1.5 µl of forward and reverse primers (2 mM each; **Extended Data Table 2**), 5 µl of SYBR® Premix Ex Taq (Tli RNase H Plus) (Takara) and run in a CFX connect instrument (Bio-Rad). PCR cycling profile consisted in a denaturing step at 95 °C for 30 s and 40 cycles at 95 °C for 10 s and 60 °C for 30 s. miR-430b mRNA was used as control.

### *In vivo* RNA integrity analysis

RNA samples containing 10 embryos and purified using standard TRIzol protocol, were submitted to RNA integrity analysis using Bioanalyzer Agilent 2100 (**Extended Data Fig. 3B, Extended Data Fig. 7A, B and G, Extended Data Fig. 7C and G, and Extended Data Fig. 9K-L**).

For *in vivo* RNA integrity analysis of IVTed gRNAs (**Fig. 2C-E and H and Extended Data Fig. 8D**), 300-500 ng of total RNA was heat denatured for 2 min at 70°C, placed on ice, and then loaded onto an RNA Nano Chip for the Agilent Bioanalyzer 2100, prepared according to manufacturer’s protocol. Electrophoresis results were exported as image representations and as raw .csv files of fluorescence intensity and time, including the ladder. Using R-Studio, the ladder was fit with a second-degree polynomial and the polynomial fit was used to calculate the size of the species in each experimental lane, resulting in a table with of traces of all RNA species represented by size in nucleotides and fluorescence intensity. Size values fit with regions of the second-degree polynomial with a negative derivative (including those of the 25 nt lower marker) were then trimmed from these traces. A custom pipeline based around findpeaks (pracma library) to detect, identify, quantify, and analyze up to the 3 most prominent RNA species (18S, 28S, long 28S cleavage product) in each trace (based on max-normalization) was developed and employed. If a species was absent based on our peak calling results, the intensity within the search window for that species was averaged and this mean intensity was used in calculating the “28S rRNA integrity ratio”. In order to obtain reliable results of 28S rRNA integrity ratio, all the samples must be run in the same gel electrophoresis within the 2100 Bioanalyzer.

### *In vitro* RNA integrity analysis

A single *in vitro* experimental rRNA integrity assay (**Extended Data Fig. 3C-D**) consists of the following components in a 10 μL reaction: 60 ng of RfxCas13d protein, 500 ng of gRNA of interest, 300-500 ng zebrafish total RNA, 1 μL of 10X CutSmart Buffer (New England Biolabs), and nuclease free water to 10 μL. Reactions were incubated at 28.5°C for 45 minutes, heat denatured for 2 min at 70°C, placed on ice, and then 1 μL of the reaction was loaded onto an RNA Nano Chip for the Agilent Bioanalyzer 2100, prepared according to manufacturer’s protocol. Electrophoresis results were exported as image representations and as raw .csv files of fluorescence intensity and time, including the ladder.

Using R-Studio, the ladder was fit with a second-degree polynomial and trimmed traces as above. These traces were further timed to remove the gRNA peak. *In vivo* rRNA analysis pipeline (see above) was modified and based around findpeaks() (pracma library) to detect, identify, quantify, and analyze up to the 3 most prominent RNA species (18S, 28S, long 28S cleavage product) in each *in vitro* trace (based on max-normalization). Due to the 10x lower input of RNA in the *in vitro* traces, short 28S cleavage product was not detected above the baseline. If a species was absent based on our peak calling results, the intensity within the search window for that species were averaged and this mean intensity was used in calculating the “28S rRNA integrity ratio”. In order to obtain reliable results of 28S rRNA integrity ratio, all the samples must be run in the same gel electrophoresis within the 2100 Bioanalyzer.

### RNA-seq libraries and analysis

Between 10 to 20 zebrafish embryos per biological replicate were collected at 4 or 6 hpf and snap-frozen. For analyzing pri-miR430 targets stability with optimized NLS version of RfxCas13d (**Fig. 3**) and to determine the collateral activity induced by different CRISPR-Cas RNA targeting systems (**Figs. 5 and 6, Extended Data Fig. 11**), total RNA was isolated using standard TRIzol protocol as described in the manufacturer’s instructions (ThermoFisher Scientific) and quantified using the Qubit fluorometric quantification (#Q10210, ThermoFisher Scientific).

For pri-miR430 targets stability, 200 ng (except for 13.5 ng and 174 ng, for two samples) of high-quality total RNA was used, as assessed using the Bioanalyzer (Agilent), with the NEBNext Poly(A) mRNA Magnetic Isolation Module (NEB, Cat. No. E7490L) at a 1/3^rd^ reaction volume. Purified mRNA was processed using the NEBNext Ultra II Directional RNA Library Prep Kit for Illumina (NEB, Cat. No. E7760L) at a 1/10^th^ reaction volume. Poly(A) isolated mRNA was resuspended in 2.25 µL fragmentation mix and fragmented for 15 min at 94°C then placed on ice for 2 min. First strand cDNA synthesis was performed by manually transferring 1 µL of the fragmented mRNA into a 384 well Armadillo PCR microplate (ThermoFisher, AB2396), containing 1 µL of the First Strand cDNA synthesis reaction master mix aliquoted by the Mosquito HV Genomics (SPT Labtech) nanoliter liquid-handling instrument. Second strand synthesis was completed per protocol at the miniaturization scale and the cDNA was purified using the SPRIselect bead-based reagent (Beckman Coulter, Cat. No. B23318) at 1.8X with the Mosquito HV and eluted in 5 µL of 0.1X TE. Libraries were generated using the NEBNext Ultra II Directional RNA Library Prep Kit at a 1/10th reaction volume starting with 5 µL of cDNA. Adaptors were ligated by adding 500 nL of NEBNext Universal Adaptor diluted at 20-fold in supplied adaptor dilution buffer. The adaptor-ligated material was PCR amplified with 14 cycles using the NEBNext Multiplex Oligos for Illumina (96 Unique Dual Index Primer Pairs) (NEB, Cat. No. E6442S) and the indexed libraries were purified using SPRIselect at 0.9X with the Mosquito HV and eluted in 10 µL of 0.1X TE.

For collateral activity RNAseq, mRNAseq libraries were generated from 100 ng (or ≤100 ng; RfxCas13d and DjCas13d) or 200 ng (or 40 ng for one sample; Cas7-11) of high-quality total RNA, and analyzed using the Bioanalyzer (Agilent). Libraries were made according to the manufacturer’s directions using a 25-fold (or 100-fold) dilution of the universal adaptor and 12-16 cycles of PCR per the respective masses with the NEBNext Ultra II Directional RNA Library Prep Kit for Illumina (NEB, Cat. No. E7760L), the NEBNext Poly(A) mRNA Magnetic Isolation Module (NEB, Cat. No. E7490L), and the NEBNext Multiplex Oligos for Illumina (96 Unique Dual Index Primer Pairs) (NEB, Cat. No. E6440S) and purified using the SPRIselect bead-based reagent (Beckman Coulter, Cat. No. B23318).

Generated short fragment libraries for **Fig. 3, 5 and 6 and Extended Data Fig. 11** were checked for quality and quantity using the Bioanalyzer and the Qubit Flex Fluorometer (Life Technologies). Equal molar libraries were pooled, quantified, and converted to process on the Singular Genomics G4 with the SG Library Compatibility Kit, following the “Adapting Libraries for the G4 – Retaining Original Indices” protocol. The converted pool was sequenced on an F3 flow cell (Cat. No. 700125) on the G4 instrument with the PP1 and PP2 custom index primers included in the SG Library Compatibility Kit (Cat. No. 700141), using Instrument Control Software 23.08.1-1 with the following read length: 8 bp Index1, 100 bp Read1, and 8 bp Index2. Following sequencing, sgdemux 1.2.0 was run to demultiplex reads for all libraries and generate FASTQ files.

To determine the number of gRNAs that can be injected together to detect highly efficient gRNAs with CRISPR-RfxCas13d (**Extended Data Fig. 5**) and then to analyze the prediction of *ex vivo* computational models using *in vivo* data from 200 gRNAs injected in sets of 25 gRNAs (**Fig. 4 and Extended Data Fig. 6**), total RNA was isolated at 4 hpf using Direct-zol RNA Miniprep Kit (#R2050, Zymo Research) following manufacturer’s instructions and quantified using the Qubit fluorometric quantification (#Q10210, ThermoFisher Scientific). cDNA was generated from 1.25 ng (2.5 ng for two replicates of RfxCas13d control samples) of high-quality total RNA, as assessed using the Bioanalyzer (Agilent), according to manufacturer’s directions for the SMART-seq v4 Ultra Low Input RNA Kit (Takara, 634891) at a 1/8^th^ reaction volume (1/4^th^ reaction volume for two replicates of RfxCas13d control) and using the Mantis (Formulatrix) nanoliter liquid-handling instrument to pipette the reagents for cDNA synthesis. Libraries were generated manually (or with the Mosquito HV Genomics (SPT Labtech) nanoliter liquid-handling instrument for RNAseq in **Fig. 4 and Extended Data Fig. 6**), using the Nextera XT DNA Library Preparation Kit (Illumina, FC-131-1096) at 1/8th reaction volumes paired with IDT for Illumina DNA/RNA UD Indexes Set A (Illumina, 20027213), and purified using the Ampure XP bead-based reagent (Beckman Coulter, Cat. No. A63882). Resulting short fragment libraries were checked for quality and quantity using the Bioanalyzer and Qubit Fluorometer (ThermoFisher). Equal molar libraries were pooled, quantified, and sequenced on a High-Output flow cell of an Illumina NextSeq 500 instrument using NextSeq Control Software 2.2.0.4 (or NextSeq Control Software 4.0.1 for RNAseq in **Fig. 4 and Extended Data Fig. 6**) with the following read length: 70 bp Read1, 10 bp i7 Index and 10 bp i5 Index. Following sequencing, Illumina Primary Analysis version NextSeq RTA 2.4.11 (or version NextSeq RTA 2.11.3.0 for RNAseq in **Fig. 4 and Extended Data Fig. 6**) and Secondary Analysis version bcl2fastq2 v2.20 were run to demultiplex reads for all libraries and generate FASTQ files.

RNA-seq reads were demultiplexed into Fastq format allowing up to one mismatch using Illumina bcl-convert 3.10.5. Reads were aligned using STAR version 2.7.3a to *Danio rerio* reference genome *danRer11* from University of California at Santa Cruz (UCSC) with GFP exogenous sequence incorporated in its index using Ensembl 106 gene models.

TPM (Transcript per Million) values were generated using RSEM version 1.3.0. Fold change for each gene was calculated using deseq2 (1.42.0) R package after filtering genes with a count of less than 10 reads in all control libraries. The resulting *p*-values were adjusted with Benjamini-Hochberg method using R function p.adjust. For collateral activity assay, genes with less than 20 counts in the control conditions were filtered.

### Guide RNAs efficacy estimation

To determine the number of gRNAs that could be injected together in zebrafish embryos to detect highly active gRNAs, we co-injected 10 and 25 gRNAs together with RfxCas13d protein in one-cell stage zebrafish embryos (**Extended Data Table 1**). Then, RNAseq at 4 hpf were performed as described earlier and gRNAs were divided into quintiles according to their activity (q5 and q1 being the most and the least efficient gRNAs, respectively, **Extended Data Fig. 5**).

To calculate gRNAs efficacy *in* vivo, 200 gRNAs (**Extended Data Table 1**) were injected in 8 different sets of 25 gRNAs together with RfxCas13d protein in one-cell stage zebrafish embryos. Two to three gRNAs targeting the same mRNA (75 mRNAs in total) were designed as described above, specifically to the longest transcript isoform. gRNAs from set 1, set 2 and set 3 targeted the same 25 mRNAs, gRNAs from set 4, set 5 and set 6 targeted the following 25 mRNAs, and gRNAs from set 7 and set 8 targeted the last 25 mRNAs. For instance, *aebp2* transcript, one of the first 25 mRNAs, was targeted with gRNA1 (13d_aebp2_1) in set 1, gRNA2 (13d_aebp2_1) in set 2 and gRNA3 (13d_aebp2_1) in set 3. Then, *in vivo* gRNA efficacy for each individual gRNA was calculated as the inverse of the Fold Change, obtained from RNAseq from zebrafish embryos at 4 hpf.

The most recent and updated computational models to predict CRISPR-RfxCas13d activity generated from *ex vivo* cell culture data, TIGER^17^, DeepCas13^18^ and RNAtargeting^19^, were used to estimate gRNA efficacy. Since RNAtargeting generates gRNA spacer sequences of 30 nt length, the first 23 nt from the 5’ end were used to make them comparable to our gRNAs and to other models. Out of 200 gRNA used in this experiment, including 3 gRNAs as negative controls, 191 were compatible to be analyzed in all computational models. Then, the performance of each prediction model was evaluated using Pearson’s correlation coefficient. *Ex vivo* cell culture data from 396 gRNAs targeting *gfp* mRNA^16^ was used as an external control.

### Statistical analyses

All statistical analyses were performed without predetermining sample size. The experiments were not randomized, and investigators were not blinded to allocation during experiments and outcome assessment. No data was excluded from the analysis. Number of embryos, replicates and experiments are indicated in figures and/or figure legends.

For phenotypes derived from embryo microinjections, Xi-square or Fisher statistical analyses were undertaken using GraphPad Prism 8 (La Jolla, CA, USA). For qRT-PCR and GFP and DsRed fluorescence levels, T-test statistical analyses were performed. *p*-values are indicated in figures or figure legends. Non-parametric Mann-Whitney U statistical tests were performed to compare high (top 5) and low (bottom 5) active gRNAs scores (**Fig. 4C-E**).

*p*-values and distances (D, maximal vertical distance between the compared distribution) for the comparison of the cumulative distribution of RNA levels at 6 hpf (**Fig. 3C**) were calculated using Kolmogorov-Smirnov Tests by dgof (v 1.4) in R package.

## Data availability

Sequencing data have been uploaded to Gene Expression Omnibus (GEO) (Series GSE270724). Imaging and raw data are available upon request. All other data is available in the main text or the supplementary information.

## Acknowledgements

We thank all members of the Moreno-Mateos laboratory for intellectual and technical support. This work was supported by Ramon y Cajal (RyC-2017-23041), PID2021-127535NB-I00, CNS2022-135564 and CEX2020-001088-M grants funded by MICIU/AEI/ 10.13039/501100011033 by “ERDF A way of making Europe” (“ERDF/EU”), and by ESF Investing in your future from Ministerio de Ciencia, Innovación y Universidades and European Union (M.A.M.-M.). This work has also been co-financed by the Spanish Ministry of Science and Innovation with funds from the European Union NextGenerationEU (PRTR-C17.I1) and the Regional Ministry of University, Research and Innovation of the Autonomous Community of Andalusia within the framework of the Biotechnology Plan applied to Health. The Moreno-Mateos lab was also funded by European Regional Development Fund (FEDER 80% of the total funding) by the Ministry of Economy, Knowledge, Business and University, of the Government of Andalusia, within the framework of the FEDER Andalusia 2014-2020 operational program within the objective “Promotion and generation of frontier knowledge and knowledge oriented to the challenges of society, development of emerging technologies (grant UPO-1380590)”. M.A.M.-M. was the recipient of the Genome Engineer Innovation 2019 Grant from Synthego. The CABD is an institution funded by University Pablo de Olavide, Consejo Superior de Investigaciones Científicas (CSIC), and Junta de Andalucía. I.M-S. was a recipient of the Margarita Salas Postdoctoral contract funded by “NextGenerationEU”, Plan de Recuperación, Transformación y Resilencia and Ministerio de Ciencia, Innovación y Universidades (recualificación del sistema universitario español 2021-2023, Pablo de Olavide University Call). L.H-H. and D.N-C. were recipients of Ayudas para contratos predoctorales para la formación de doctores (Ministerio de Ciencia e Innovación) funded by MICIU/AEI /10.13039/501100011033 and FSE invierte en tu futuro and FSE. C.G-M. was funded with Ayudas captación, incorporación y movilidad de capital humano de I+D+i, Junta de Andalucía (POSTDOC 21_00667). P.M.M.G. was funded by a postdoctoral fellowship from Junta de Andalucía (DOC_00397). This study was supported by the Stowers Institute for Medical Research. A.A.B. was awarded a US National Institutes of Health grant (NIH-R01 GM136849 and NIH R21OD034161). A.J.T. was supported by the US National Institute of Health: F31HD110268. This work was completed as part of thesis research for A.J.T. & G.dS.P., Graduate School of the Stowers Institute for Medical Research. This work was supported by grant PID2021-125682NB-I00 to M.A.N. funded by MICIU/AEI/10.13039/501100011033 and by FEDER, UE, and by Instituto de Salud Carlos III (CIBERER, CB19/07/00038 to MAN), who also acknowledges financial support from the Spanish State Research Agency, through the “Severo Ochoa Program” for Centres of Excellence in R&D Grant CEX2021-001165-S funded by MCIN/AEI/ 10.13039/501100011033. M.J.M. was supported by grant PID2020-120463RB-I00 funded by the Spanish Ministerio de Ciencia e Innovación. We thank our colleague Francisco J. Guerra (Stowers Institute and CABD) for the initial research on Cas7-11. We also thank Rhonda Egidy and Anoja Perera from Sequencing and Discovery Genomics at Stowers (Kansas, MO, USA).

## Author contributions

M.A.M-M. conceived the project and designed the research. I.M-S., L.H-H. and D.N-C. performed most of the experiments. C.G-M., A.C-R. and M.J.M. contributed in the CRISPR-Cas7-11 experiments and analysis. A.J.T., G.dS.P. and G.K. performed part of the experiments and analyzed the results from *in vitro* transcribed gRNAs assays. L.T-G. and A.D-M. performed protein purifications. J.A.W.II produced the synthetic gRNAs. K.H helped with cm-gRNAs section together with J.G. and M.A.N. that also contributed with the optimization of RfxCas13d-NLS. P.M.M-G. performed gRNA activity prediction analysis and computational models’ comparison. M.A.M-M. I.M-S., L.H-H. and D.N-C. carried out data analysis with the help of A.A.B. M.A.M-M. and I.M-S. wrote the manuscript with the contribution of L.H-H., D.N-C. and A.A.B. and with the input from the other authors. All authors reviewed and approved the manuscript.

