## Supplementary figures for "Enhanced RNA-targeting CRISPR-Cas technology in zebrafish"

#### Extended Data Figures

**Extended Data Figure 1. RfxCas13d protein is less efficient than *RfxCas13d* mRNA when targeting mid- and late-zygotically expressed mRNAs.**

**A)** Representative images of the phenotypes obtained after the injection of the mRNA-gRNAs or ribonucleoprotein (RNP) complexes targeting zygotically-expressed *slc45a2 – albino*, and *slc24a5 – golden*. Loss-of-function phenotypes (reduced or lack of pigmentation) were evaluated at 48 hpf (scale bar, 0.75 mm for lateral views and 0.1 mm for insets). Mild (least extreme), Severe (medium level), and Albino-like (most extreme).

**B)** Stacked barplots showing percentage of observed phenotypes in injected embryos with standard or chemically modified gRNAs (cm-gRNAs) targeting *noto*, *rx3*, *tyrosinase* and *albino* (gNOTO, gRX3, gTYR and gALB, respectively) together with RfxCas13d protein (pRfxCas13d). (n) total number of embryos is displayed for each condition. The results are shown as the averages ± standard deviation of the mean of each phenotypic category from at least two independent experiments. $\chi^2$ (for *noto*) or Fisher statistical tests were performed to compare standard and cm-gRNAs.

**C)** Stacked barplots showing percentage of observed phenotypes in injected embryos with standard or cm-gRNAs targeting *no-tail*, *noto*, *rx3*, *albino* and *golden* (1 ng of a mix of three gRNAs – gNTL, gNOTO, gALB and gGOLDEN; 1 ng of single gRNAs – gRX3-2 and gRX3-3) together with RfxCas13d mRNA (300 pg of mRfxCas13d). (n) total number of embryos is displayed for each condition. The results are shown as the averages ± standard deviation of the mean of each phenotypic category. $\chi^2$ or Fisher (for *golden*) statistical tests were performed to compare standard and cm-gRNAs.

**D)** Barplots representing mRNA relative levels analyzed by qRT-PCR at 6 hpf (*no-tail* and *noto*) or 20 hpf (*rx3* and *tyrosinase*) from injected embryos with mRfxCas13d together with gNTL, gNOTO, gRX3-1 or gTYR. Results are shown as the averages ± standard deviation of the mean from at least four biological replicates from two independent experiments. T-test statistical analyses were performed. *taf15* (for *no-tail* and *noto*) or *ef1α* (for *rx3* and *tyrosinase*) mRNA was used as a normalization control.

**Extended Data Figure 2. Increasing the length of gRNA spacers does not improve CRISPR-RfxCas13d activity in zebrafish embryos.**

**A)** Schematic illustration of the experimental setup used to compare 23 or 30 nt spacers for CRISPR-RfxCas13d gRNAs. 300 pg of RfxCas13d mRNA (mRfxCas13d) or 3 ng of RfxCas13d protein (pRfxCas13d) were injected together with 1 ng of a mix of three gRNAs (~300 pg from each gRNA) into one-cell stage zebrafish embryos.

**B, C)** Stacked barplots showing percentage of observed phenotypes using mRfxCas13d (**B**) or pRfxCas13d (**C**) targeting *nanog*, *no-tail* and *tyrosinase* (gNANOG, gNTL, and gTYR, respectively). (n) total number of embryos is indicated for each condition. The results are shown as the averages ± standard deviation of the mean of each phenotypic category. Representative images of phenotypes are displayed in **Fig. 1C**. Fisher statistical tests were performed to compare 23nt- versus 30nt-gRNAs.

**Extended Data Figure 3. *In vitro* transcribed gRNAs can induce toxicity**

**A)** Representative images of RfxCas13d-mediated *si:dkey-93m18.4* KD and control embryos at 2, 4, and 6 hpf. Top single embryos show expected stage. Red boxes denote abnormal development (left). Representative images of developmental stages and dead embryos evaluated at ~6 hpf: sphere/high, 30% – 50% epiboly, and germ ring/shield stages correspond to 3.3 – 4, 4.6 – 5.3, 5.7 – 6 hpf in non-injected embryos growing in standard conditions, respectively (scale bar, 0.5 mm) (right).

**B)** RNA integrity images from bioanalyzer of total RNA from RfxCas13d mediated *si:dkey-93m18.4* and *brd4* KD and control embryos at 4 hpf. Red arrows denote bands from 28S rRNA cleavage.

**C)** Scheme of *in vitro* rRNA integrity assay (see Methods for further details).

**D)** RNA integrity images from at least two replicates of bioanalyzer electrophoresis of the *in vitro* rRNA integrity assay. Red arrows denote bands from 28S rRNA cleavage. Negative controls indicate *in vitro* rRNA integrity assays without RfxCas13d.

**Extended Data Figure 4. Optimization of NLS versions to increase CRISPR-RfxCas13d nuclear RNA-targeting**

**A)** Diagram illustrating the cytoplasmic (pink) and five different NLS versions of RfxCas13d: NLS (SV40 NLS, blue), 2C-NLS or optimized NLS (SV40 and nucleoplasmin long-NLP NLS, orange; Liu et al., 2019[42]), hei-tag (oNLS, green; Thumberger et al., 2022[45]), cmyc-2C-NLS (cmyc, SV40 and nucleoplasmin long-NLP NLS, purple; Wu et al., 2019[43]) and bpNLS (bipartite NLS, red; Liang et al., 2022[44]).

**B)** Stacked barplots showing percentage of observed phenotypes at 48 hpf (**Fig. 3A**) when injecting each mRNA alone (300 pg/embryo). (n) total number of embryos is indicated for each condition. The results are shown as the averages ± standard deviation of the mean of each developmental stage from at least two independent experiments.

**Extended Data Figure 5. Highly efficient CRISPR-RfxCas13d gRNAs can be identified when injecting up to 25 gRNAs together.**

**A)** Schematic representation of multiplexing with CRISPR-RfxCas13d using 10 gRNAs (100 pg each gRNA – purple) or 25 gRNAs (40 pg each gRNA – orange).

**B)** Scatterplot showing mRNA level (RNA-seq) of zebrafish embryos injected with RfxCas13d protein (3 ng/embryo) together with 10 or 25 gRNAs compared with embryos injected only with RfxCas13d protein at 4 hpf. gRNAs are color-coded in quintiles (q). First 10 targeted mRNAs are depicted in circles and 15 mRNAs up to 25 are indicated in squares in their respective panels.

**C)** Boxplots showing the expression levels of polyadenylated RNAs along the first 4 hours of zebrafish embryo development (log2 TPM: Transcript per million). The

75 mRNAs targeted in **Fig. 4B** and **Extended Data Fig. 6** are depicted in green. Data from Medina-Muñoz et al., 2021[49].

**Extended Data Figure 6. *In vivo* activity of CRISPR-RfxCas13d gRNAs activity predicted by *ex vivo* computational models.**

**A)** Volcano plots showing mRNAs targeted (red) with each set of 25 gRNAs co-injected with RfxCas13d protein in one-cell stage embryos and measured at 4 hpf by RNAseq.

**B)** RNA integrity images from Bioanalyzer analysis of the indicated samples. RNA samples purified at 4 hpf from injected embryos with each set of 25 gRNAs and RfxCas13d protein.

**C)** Heatmap of *p*-values obtained from Pearson's correlations performed with each gRNA set and prediction scores by TIGER (Wessels et al., 2023[17]), DeepCas13 (Cheng et al., 2023[18]) and RNAtargeting (Wei et al., 2023[19]) represented in **Fig. 4B**. Correlation with cell culture data from *gfp* mRNA from 396 gRNAs (Wessels et al., 2020[16]) was used as a control.

**Extended Data Figure 7. Highly abundant *dsRed* mRNA partially buffers the collateral activity triggered by CRISPR-RfxCas13d.**

**A, B)** RNA integrity images from Bioanalyzer analysis of the indicated samples. RNA samples purified at 6 hpf from injected embryos with 300 pg RfxCas13d mRNA (mRfxCas13d, **A**) or 3 ng RfxCas13d protein (pRfxCas13d, **B**) together with 1 ng of a mix of three gRNAs targeting *gfp* (gGFP) and 10, 20, 50 or 100 pg of ectopic *gfp* mRNA (mGFP). Red arrowheads indicate major and minor bands of degradation of 28S ribosomal subunit.

**C)** Dot-blot displaying RNA integrity number calculated by Bioanalyzer from indicated samples shown in **A** and **B**. Results are shown as the averages ± standard deviation of the mean from at least two biological replicates. Data from samples injected with mRfxCas13d or pRfxCas13d are depicted in blue or orange, respectively.

**D)** Stacked barplots representing developmental phenotypes (epiboly stages) of injected embryos with pRfxCas13d together with gGFP, 75 pg of *dsRed* mRNA (mDsRed) and 10, 20 or 50 pg of mGFP. Representative images of indicated epiboly stages are shown in **Fig. 1C**. (n) total number of embryos is indicated for each condition. The results are shown as the averages ± standard deviation of the mean of each developmental stage from at least two independent experiments.

**E, F)** Barplots displaying GFP (**E**) and DsRed (**F**) fluorescence levels of injected embryos with pRfxCas13d together with gGFP, mDsRed and 10, 20 or 50 pg of mGFP. Results are shown as the averages ± standard deviation of the mean from three biological replicates of 5 embryos each. T-test statistical analyses were performed, *p*-value is indicated above.

**G)** RNA integrity analysis of the indicated samples by Bioanalyzer. RNA samples purified at 6 hpf from injected embryos with pRfxCas13d together with gGFP, mDsRed and 10, 20 or 50 pg of mGFP. Red arrowhead indicates degradation of 28S ribosomal subunit.

**H)** Volcano plot showing down- (purple) and upregulated (orange) genes upon *gfp* mRNA KD (10 or 50 pg) with pRfxCas13d. Fold-Change of 3 and $p$-value $< 0.001$ (dashed lines) were set to determine deregulated genes.

**Extended Data Figure 8. CRISPR-RfxCas13d does not induce a physiologically relevant collateral activity when targeting highly expressed endogenous mRNAs.**

**A)** Boxplots showing the expression levels of all polyadenylated RNAs along the first 4 hours of zebrafish embryo development. *hnrnpa0l*, *hmga1a* and *hspa8* mRNAs are depicted in purple, green and blue, respectively. Data from Medina-Muñoz et al., 2021[49].

**B)** Barplots exhibiting GFP fluorescence relative levels analyzed of injected embryos with pRfxCas13d together with gRNAs targeting endogenous *hnrnpa0l*, *hmga1a* and *hspa8* transcripts (gHNRNPA0L, gHMGA1A and gHSPA8, respectively) and 50 pg of *gfp* (mGFP) as a collateral activity reporter control. Results are shown as the averages ± standard deviation of the mean from six biological replicates of 5 embryos from two independent experiments. T-test statistical analyses were performed, *p*-values are indicated above.

**C)** RNA integrity analysis by Bioanalyzer of RNA samples at 6 hpf from injected embryos with pRfxCas13d together with gHNRNPA0L, gHMGA1A or gHSPA8 and 50 pg mGFP.

**D)** Boxplot of 28S integrity ratio *in vivo* at 6 hpf from injected embryos with pRfxCas13d together with gHNRNPA0L, gHMGA1A or gHSPA8 and 50 pg mGFP. Two biological replicates were analyzed. The mean, first and third quartile are represented for each condition. One-way ANOVA followed by Dunnett's post-hoc analysis was performed.

**E)** Stacked barplots representing developmental phenotypes (epiboly stages) of injected embryos with pRfxCas13d together with gHNRNPA0L, gHMGA1A or gHSPA8. Representative images of indicated epiboly stages are shown in **Fig. 1C**. $X^2$ statistical analyses were performed comparing side-by-side each knockdown condition with embryos injected only with pRfxCas13d (*p*-value indicated above). (n) total number of embryos is indicated for each condition. The results are shown as the averages ± standard deviation of the mean of each developmental stage from at least two independent experiments.

**F)** Barplots indicating mRNA relative levels analyzed by qRT-PCR of injected embryos with pRfxCas13d together with gHNRNPA0L, gHMGA1A or gHSPA8. T-test statistical analyses were performed comparing side-by-side each KD condition with embryos injected only with pRfxCas13d. Results are shown as the averages ± standard deviation of the mean from four biological replicates from two independent experiments. *taf15* mRNA was used as a normalization control.

**G)** RNA integrity analysis of the *hnrnpa0l*, *hmga1a* and *hspa8* mRNA KD by Bioanalyzer. RNA samples purified at 6 hpf from injected embryos with pRfxCas13d together with gHNRNPA0L, gHMGA1A or gHSPA8.

**Extended Data Figure 9. CRISPR-Cas7-11 RNP complexes do not exhibit collateral activity.**

**A)** Barplots showing viability (left) and toxicity (right) at 24 hpf from embryos injected with RfxCas13d, Hf-RfxCas13d, DjCas13d and Cas7-11 proteins, and RfxCas13d and Hf-RfxCas13d mRNAs. 3 ng of each purified protein were injected in one-cell stage zebrafish embryos. 1x indicates 300 pg of mRNA per embryo. (n) total number of embryos is indicated for each condition. The results are shown as the averages ± standard deviation of the mean of each developmental stage from at least three independent experiments.

**B, C)** Stacked barplots showing the percentage of observed phenotypes of injected embryos with RfxCas13d or Hf-RfxCas13d mRNA together with 1 ng of a mix of three gRNAs targeting *nanog* (**A**, gNANOG1-3) and 1 ng of one gRNA targeting *no-tail* (**B**, gNTL-4). 1x indicates 300 pg of mRNA or gRNA per embryo. Representative images of epiboly stages and *no-tail* phenotypes are exhibited in **Fig. 1C**. (n) total number of embryos is indicated for each condition. The results are shown as the averages ± standard deviation of the mean of each phenotypic category from at least two independent experiments. $X^2$ statistical tests were performed (*p*-values are indicated).

**D)** Western blot of RfxCas13d and Hf-RfxCas13d protein samples from 6 hpf zebrafish embryos injected with 300 pg of each mRNA (detected by $\alpha$-HA primary antibody). Stain-free blot image is shown as loading control.

**E, F)** Stacked barplots representing developmental phenotypes (epiboly stages) of injected embryos with DjCas13d (**E**) or Cas7-11 (**F**) together with 1 ng of a mix of three gRNAs (gGFP) targeting 10, 20, 50 or 100 pg of ectopic *gfp* mRNA (mGFP). Representative images of indicated epiboly stages are shown in **Fig. 1C**. (n) total number of embryos is indicated for each condition. The results are shown as the averages ± standard deviation of the mean of each phenotypic category from at least two independent experiments.

**G, H)** Barplots showing *gfp* mRNA relative levels by qPCR of injected embryos with DjCas13d (**G**) or Cas7-11 (**H**) together with gGFP targeting 10, 20, 50 or 100 pg of mGFP. Results are shown as the averages ± standard deviation of the mean from four biological replicates from two independent experiments. T-test statistical analyses were performed, *p*-value is indicated above.

**I, J)** Barplots showing GFP fluorescence levels of injected embryos with DjCas13d (**I**) or Cas7-11 (**J**) together with gGFP targeting 10, 20, 50 or 100 pg of mGFP. Results are shown as the averages ± standard deviation of the mean from four biological replicates from two independent experiments from at least 3 biological replicates of 5 embryos each. T-test statistical analyses were performed, *p*-value is indicated above.

**K, L)** RNA integrity analysis of the indicated samples by Bioanalyzer. RNA samples purified at 6 hpf from injected embryos with DjCas13d (**K**) or Cas7-11 (**L**) together with gGFP and 10, 20, 50 or 100 pg of mGFP.

**Extended Data Figure 10. CRISPR-DjCas13d RNP complexes induce less unspecific RNA degradation and collateral activity than CRISPR-RfxCas13d.**

**A, D)** Stacked barplots representing developmental phenotypes (epiboly stages) of injected embryos with DjCas13d (**A**) or Cas7-11 (**D**) together with gGFP, 50 pg of mGFP and 75 pg of ectopic *dsRed* mRNA (mDsRed). Representative images of indicated epiboly stages are shown in **Fig. 1C**. (n) total number of embryos is indicated for each condition. The results are shown as the averages ± standard deviation of the mean of each developmental stage from at least two independent experiments.

**B, E)** Barplots showing GFP fluorescence levels of injected embryos with DjCas13d (**B**) or Cas7-11 (**E**) together with gGFP, 50 pg of mGFP and 75 pg of mDsRed. The results are shown as the averages ± standard deviation of the mean of each developmental stage from six biological replicates of 5 embryos each. T-test statistical analyses were performed, *p*-value is indicated above.

**C, F)** Barplots showing DsRed fluorescence levels of injected embryos with DjCas13d (**C**) or Cas7-11 (**F**) together with gGFP, 50 pg of mGFP and 75 pg of mDsRed. The results are shown as the averages ± standard deviation of the mean of each developmental stage from six biological replicates of 5 embryos each. T-test statistical analyses were performed, *p*-value is indicated above.

**Extended Data Figure 11. DjCas13d did not lead to massive deregulation upon collateral activity.**

**A, B, C)** Volcano plots showing down- (purple) and upregulated (orange) genes upon 10 or 50 pg mGFP KD with RfxCas13d (**A**, same data from **Extended Data Fig. 7H**), DjCas13d (**B**) and Cas7-11 (**C**) measured by RNAseq. *gfp* levels are depicted in green. Fold-Change of 2 and *p*-value < 0.001 (dashed lines) were set to determine deregulated genes.

Moreno-Sánchez, Hernández-Huertas, Nahón-Cano et al. Extended Data Fig. 1

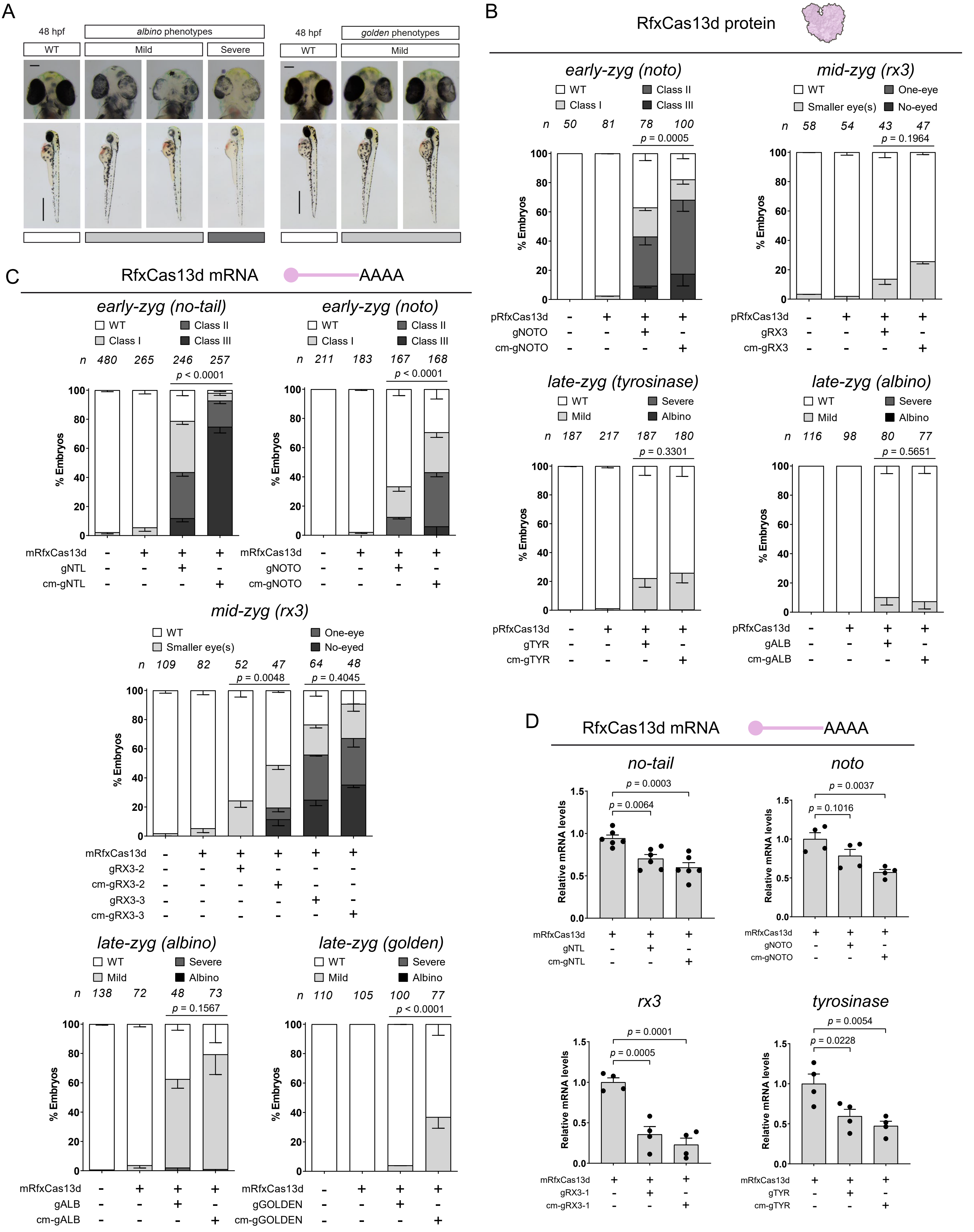

Moreno-Sánchez, Hernández-Huertas, Nahón-Cano et al. Extended Data Fig. 2

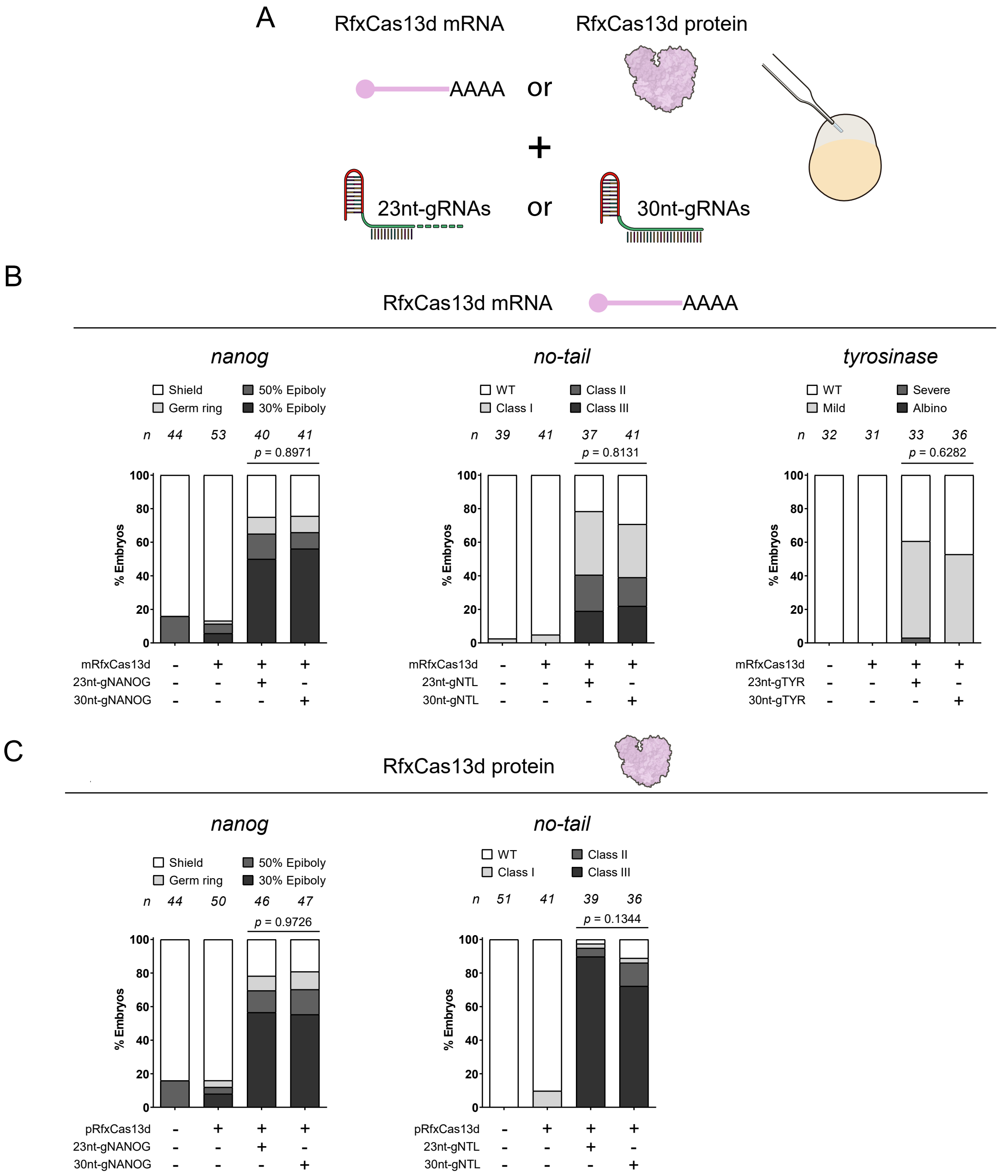

Moreno-Sánchez, Hernández-Huertas, Nahón-Cano et al. Extended Data Fig. 3

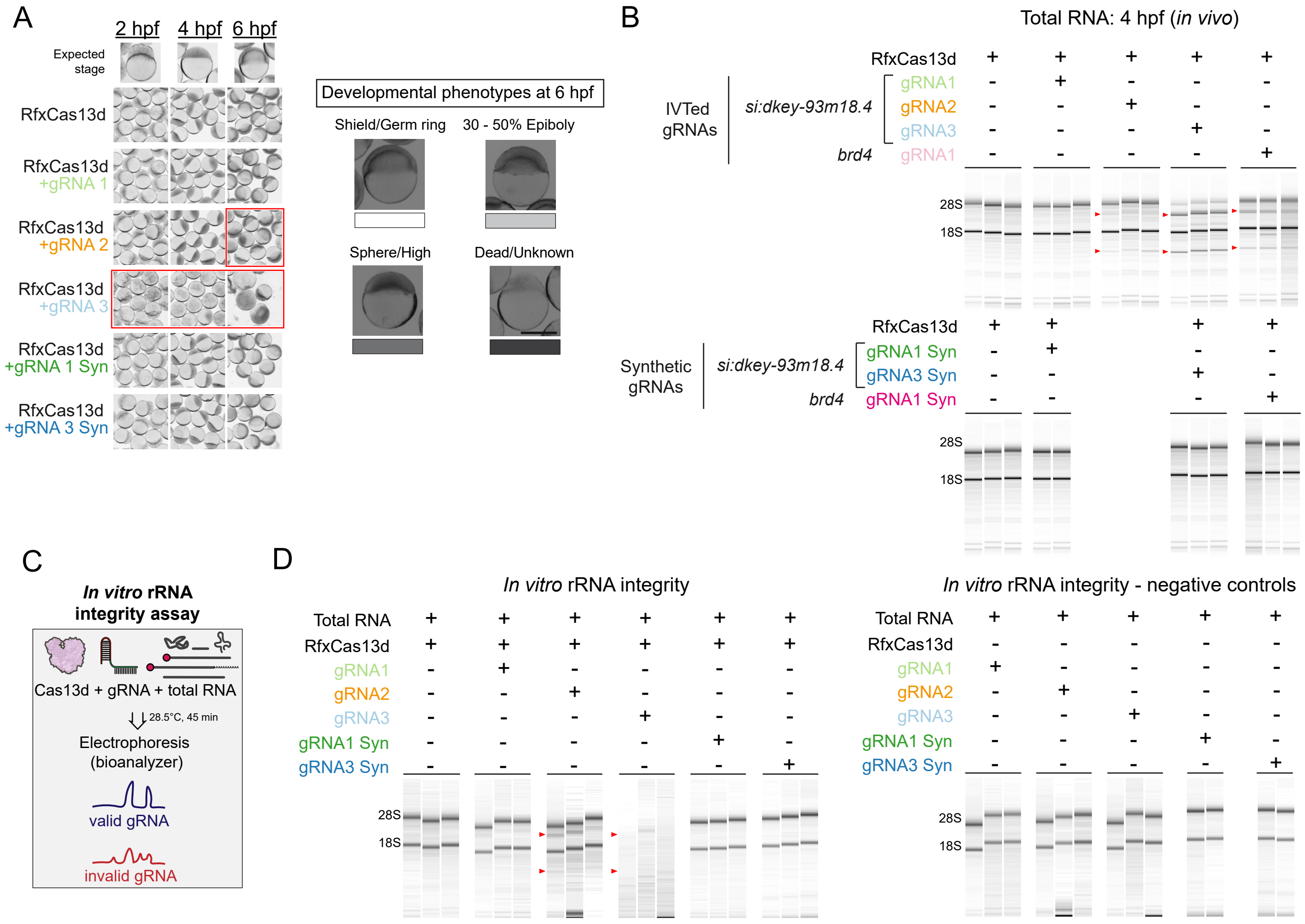

### Moreno-Sánchez, Hernández-Huertas, Nahón-Cano et al. Extended Data Fig. 4

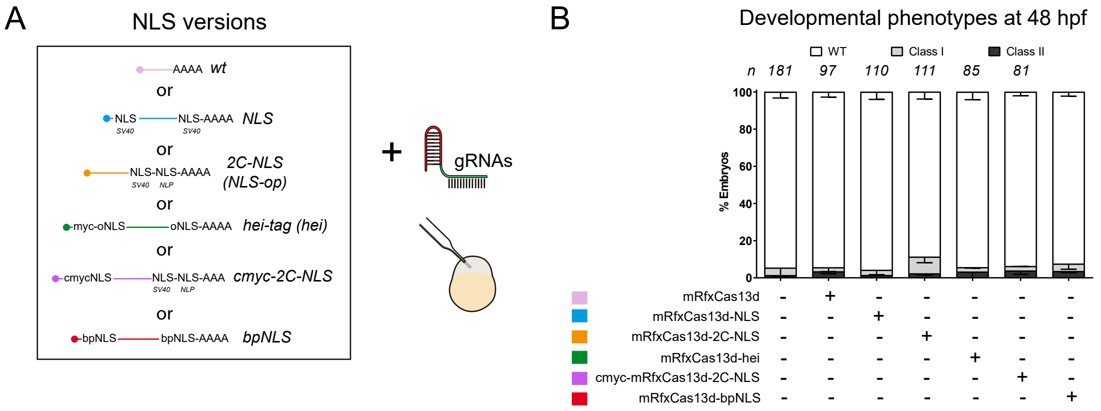

Moreno-Sánchez, Hernández-Huertas, Nahón-Cano et al. Extended Data Fig. 5

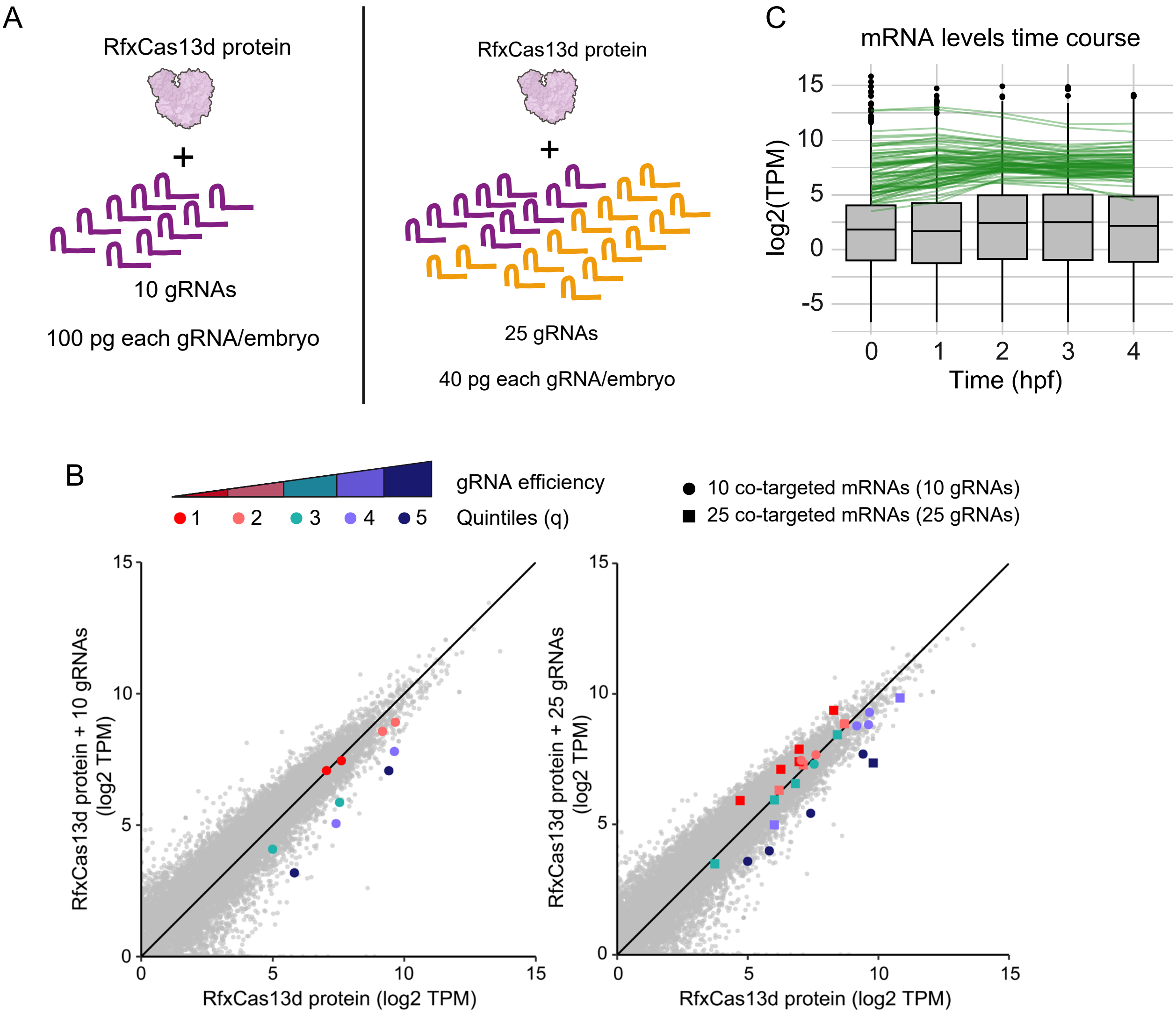

### Moreno-Sánchez, Hernández-Huertas, Nahón-Cano et al. Extended Data Fig. 6

A

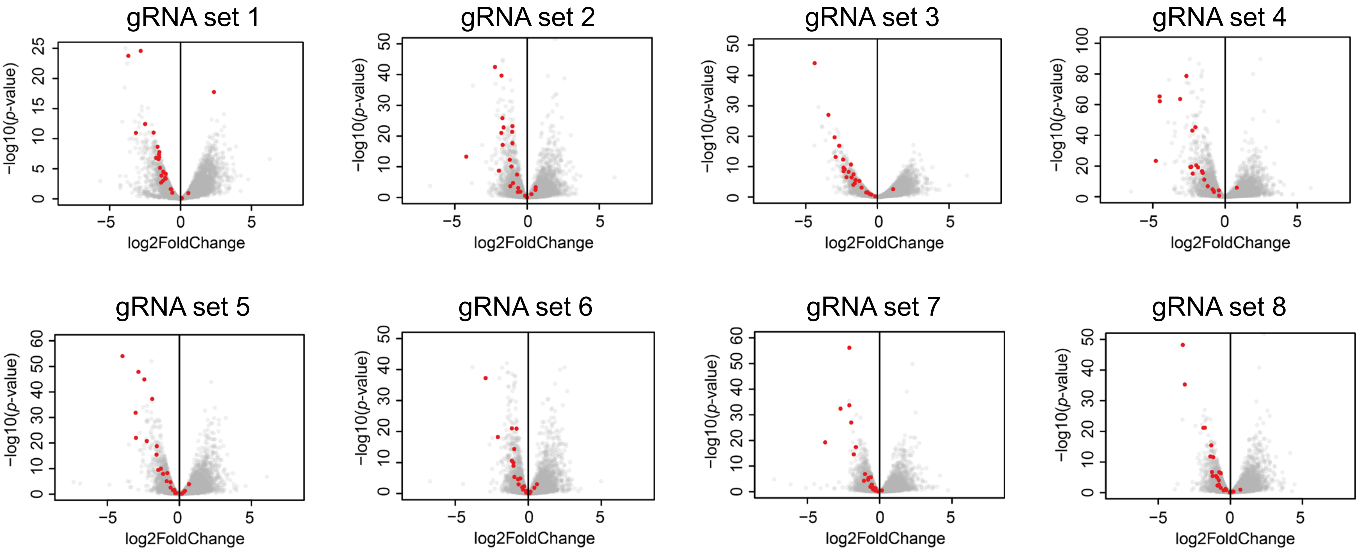


B

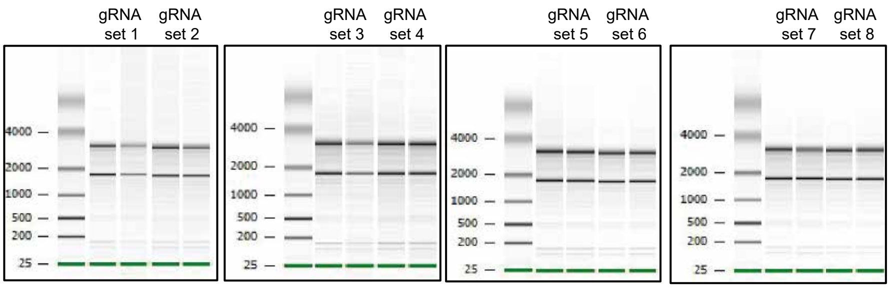


C *p*-value Pearson's correlations

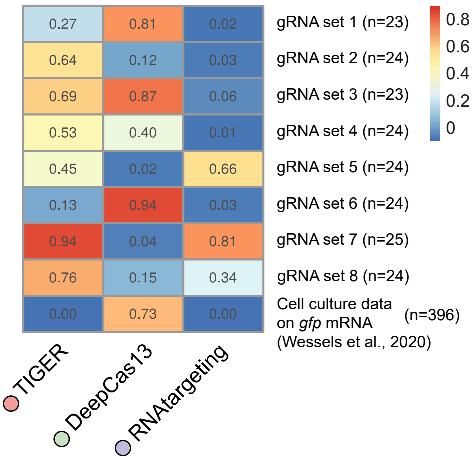

Moreno-Sánchez, Hernández-Huertas, Nahón-Cano et al. Extended Data Fig. 7

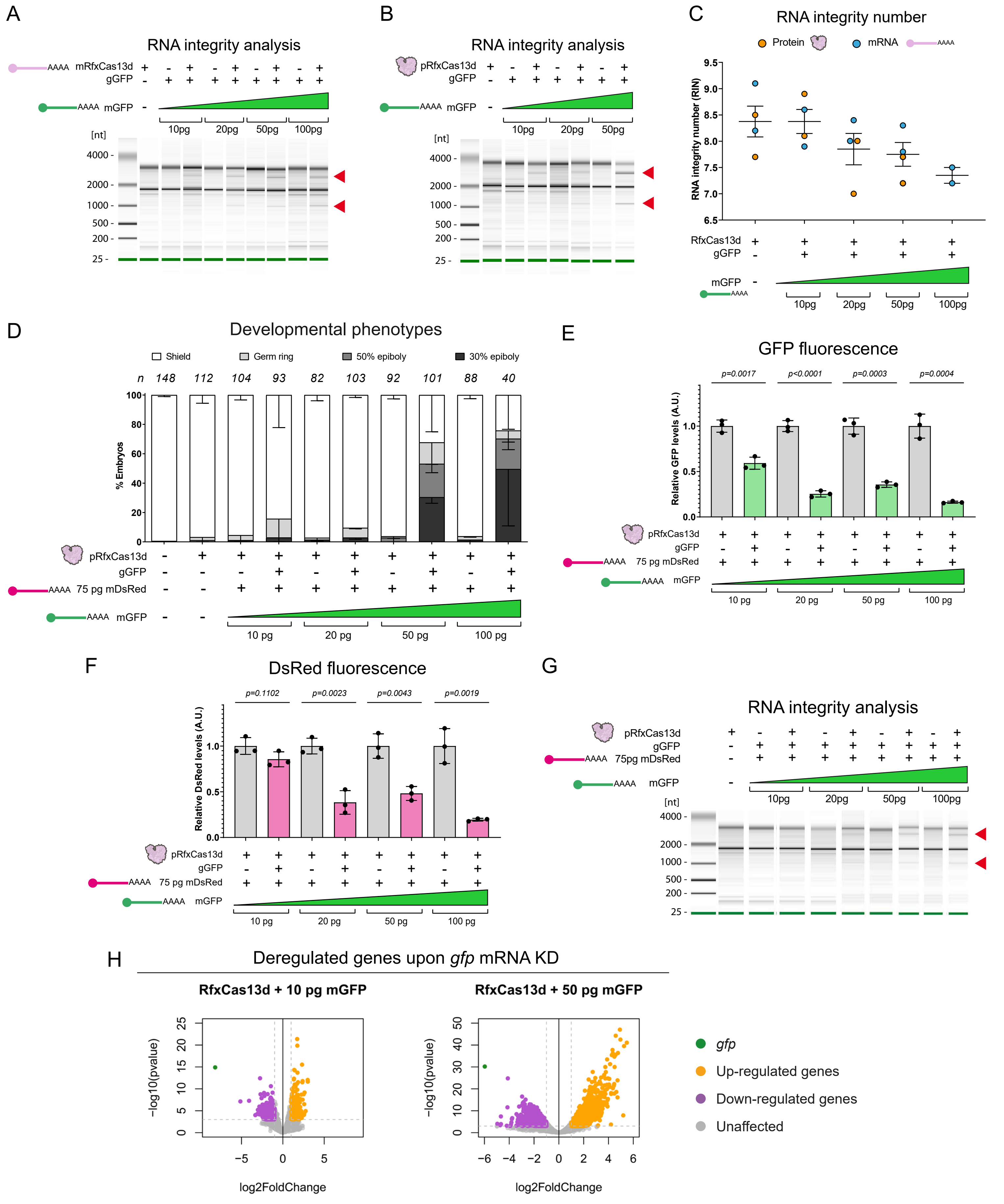

Moreno-Sánchez, Hernández-Huertas, Nahón-Cano et al. Extended Data Fig. 8

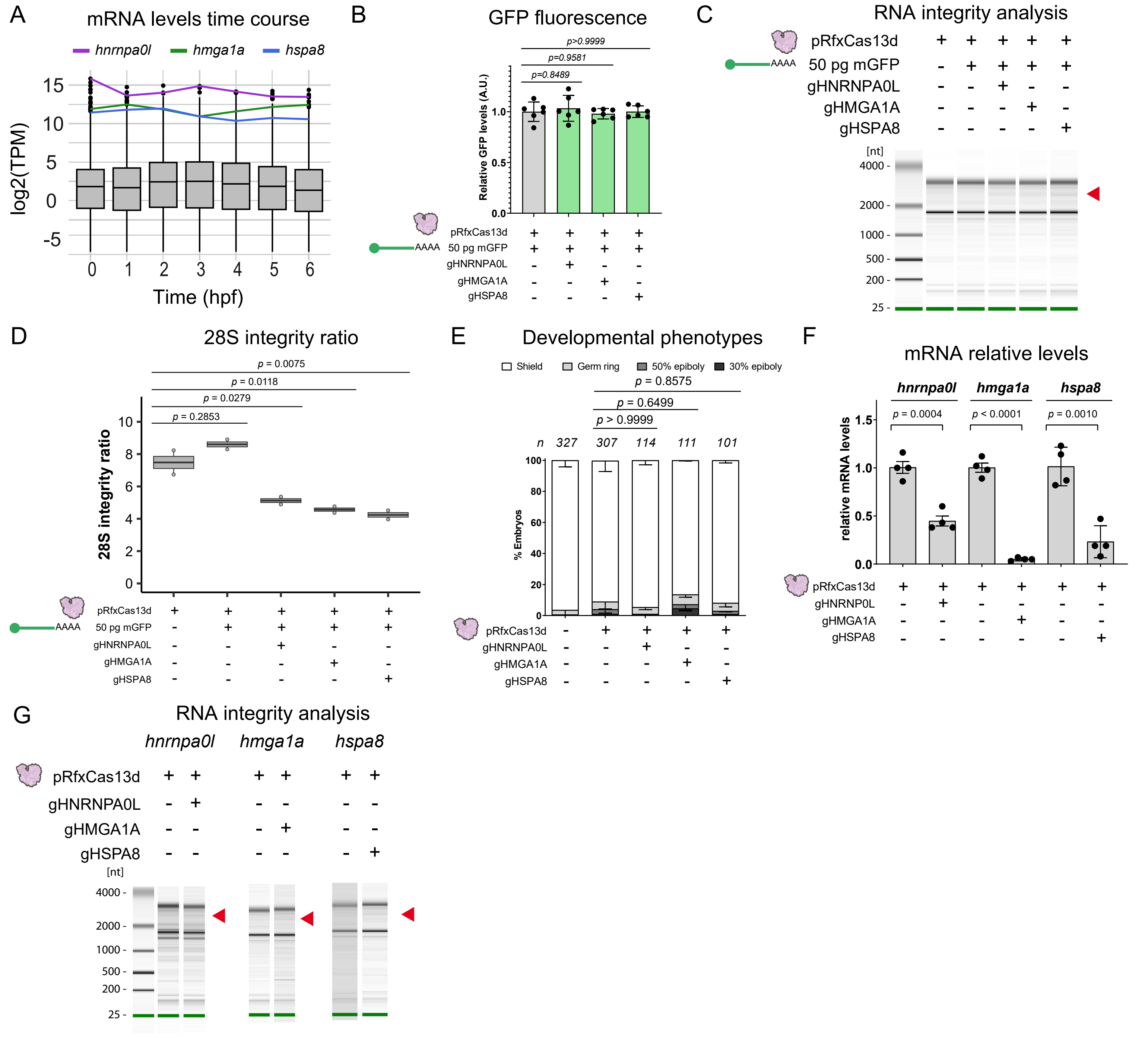

Moreno-Sánchez, Hernández-Huertas, Nahón-Cano et al. Extended Data Fig. 9

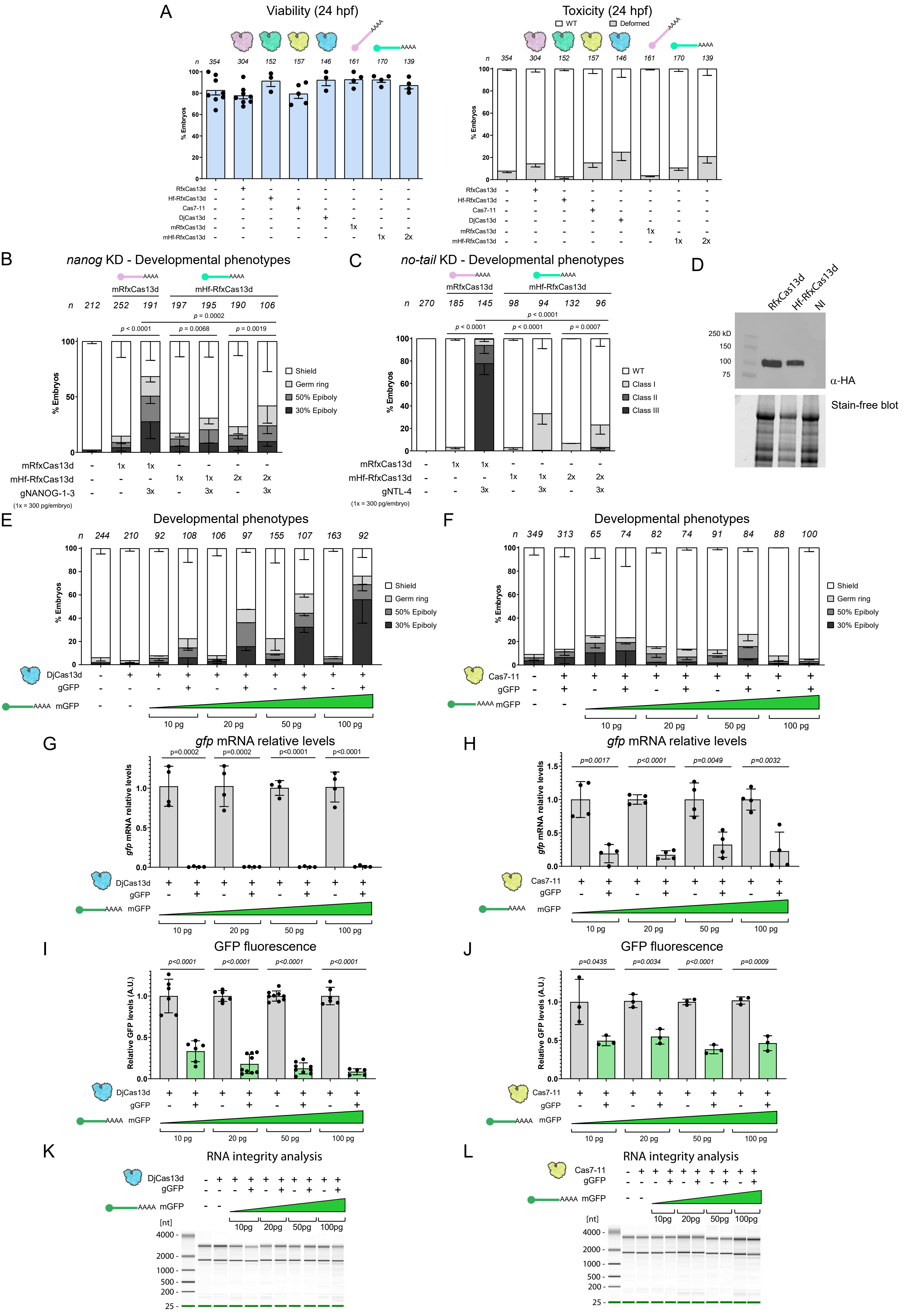

Moreno-Sánchez, Hernández-Huertas, Nahón-Cano et al. Extended Data Fig. 10

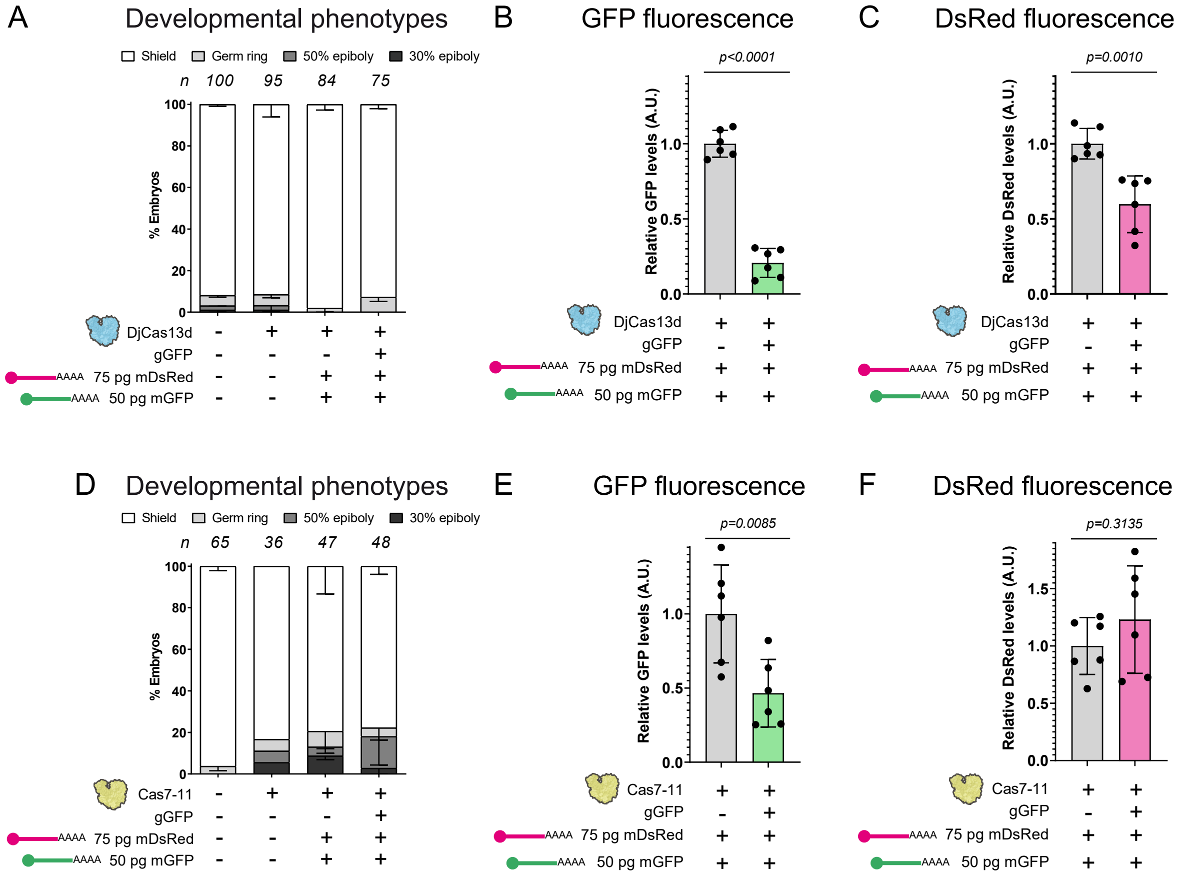

Moreno-Sánchez, Hernández-Huertas, Nahón-Cano et al. Extended Data Fig. 11

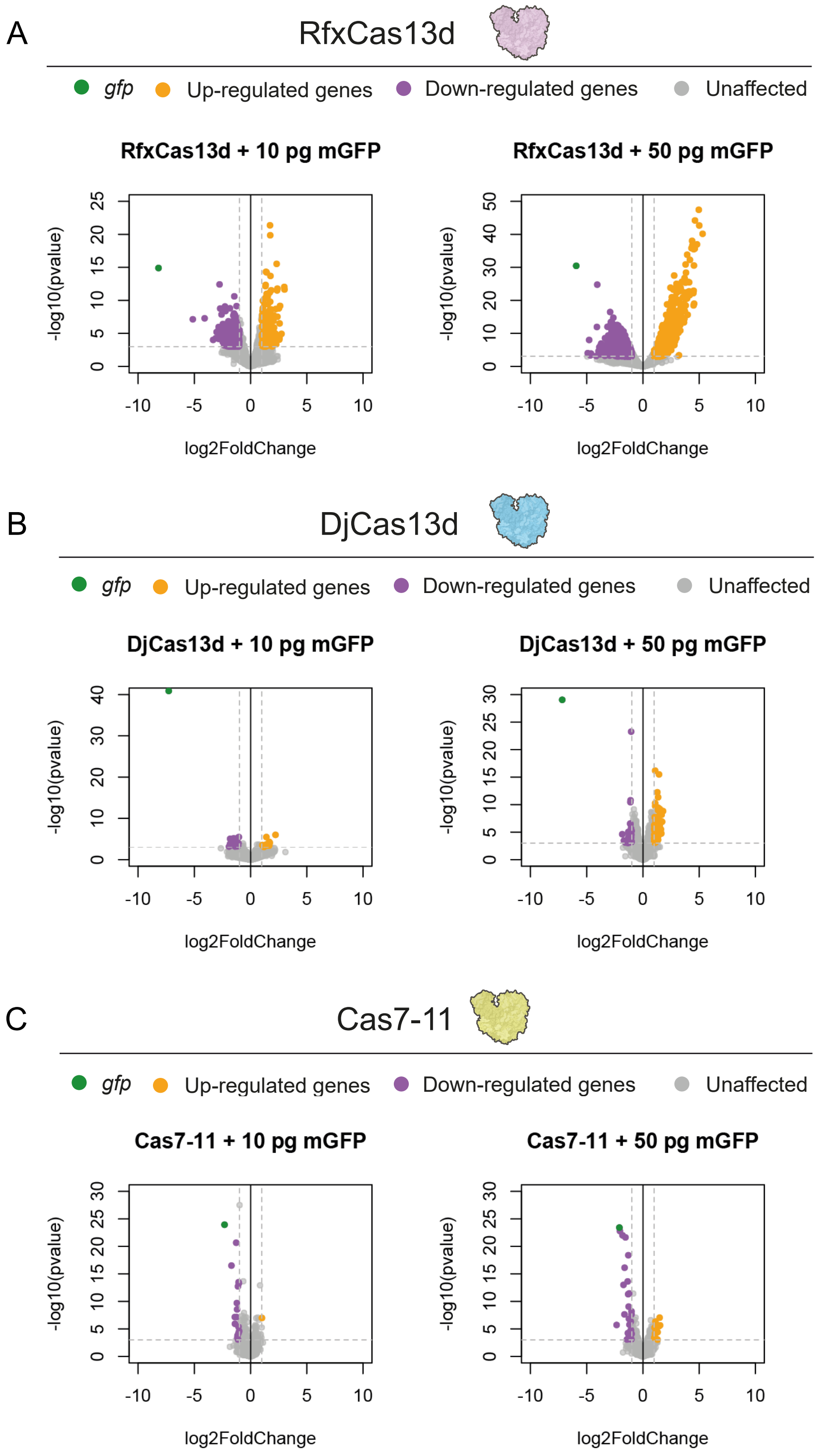
